# Snake venom-inspired novel peptides protect *Caenorhabditis elegans* against paraquat-induced Parkinson’s pathology

**DOI:** 10.1101/2024.06.01.596942

**Authors:** Dev Madhubala, Kangkon Saikia, Aparup Patra, Rosy Mahato, Pedro Alexandrino Fernandes, Arun Kumar, Mojibur R. Khan, Ashis K. Mukherjee

## Abstract

The *in vivo* protective mechanisms of two low molecular mass (∼1.4 kDa) novel custom peptides (CPs) against paraquat (PT)-induced neurodegenerative dysfunction in the *Caenorhabditis elegans* model were deciphered. CPs prevent the PT binding to the nerve ring adjacent to the pharynx in *C. elegans* (N2 strain) by stable and high-affinity binding to the tyrosine-protein kinase receptor CAM-1, resulting in significant inhibition of PT-induced toxicity by reducing enhanced reactive oxygen species production, mitochondrial membrane depolarization, and chemosensory dysfunction. The CPs inhibited PT-induced dopaminergic (DAergic) neuron degeneration and alpha-synuclein aggregation, the hallmarks of Parkinson’s Disease, in transgenic BZ555 and NL5901 strains of *C. elegans*. The transcriptomic, functional proteomics, and quantitative reverse transcription-polymerase chain reaction (qRT-PCR) analyses show that CPs prevented the increased expression of the genes involved in the skn-1 downstream pathway, thereby restoring PT-mediated oxidative stress, apoptosis, and neuronal damage in *C*. *elegans*. The CPs ability to repair PT-induced damage was demonstrated by a network of gene expression profiles illustrating the molecular relationships between the regulatory proteins. Further, CPs (10 mg/kg, parental route) did not show toxicity or induce inflammatory mediators in the mouse model.

## 1. Introduction

Parkinson’s disease (PD) is one of the second most common Neurodegenerative disorders (NDs) associated with the loss of dopaminergic (DAergic) neurons and the aggregation of alpha-synuclein (α-synuclein) [1, 2]. Paraquat (PT) is a toxic nonselective herbicide widely used for weeding in wheat fields worldwide, and agriculture workers inhale poisonous chemicals. Notably, PD risk is six times higher in agriculture workers exposed to PT than in non-exposed individuals [3]. Current therapeutic and treatment options such as levodopa or DAergic agonists (pramipexole, rotigotine, and ropinirole) for PD primarily target to increase the neurotransmitter dopamine (DA) level to improve the disease’s motor symptoms [4, 5]. However, long-term PD medication use reduces efficacy and leads to other adverse effects, such as motor complications [2, 5]. Thus, there is an urgent demand to discover alternative novel drugs for improving the management of PD.

Neurotrophins (NTs) are a subgroup of growth factors that play a role in the growth, development, protection, and repair of neurons, which are essential for the therapy of NDs [6, 7]. Nerve growth factor (NGF), one of the NTs, has been investigated for its action against NDs; however, this NT has failed in clinical trials [8, 9]. Therefore, developing novel therapeutic molecules with better potential may pave the way for new drug development against PD[10].

Snake venom is a cocktail of therapeutically active proteins and peptides [11–13]. Snake venom contains a small amount of low molecular mass protein NGF (13-17 kDa) in monomeric and dimeric forms [14–16]. Their small size, target sensitivity, and structural stability make them more advantageous over NTs as therapeutic agents for treating and preventing NDs [14, 15, 17–19]. However, purifying and characterizing proteins and peptides from snake venom is tedious and non-economical. Therefore, we designed two low molecular mass custom peptides (HNP and TNP; 1.2 KDa and 1.4 KDa) that mimic the tropomyosin receptor kinase A (TrkA) binding region of snake venom NGFs [20]. Like parent NGF molecules, the synthetic custom peptides also demonstrated *in vitro* neuritogenesis potency by binding to the TrkA receptor of rat pheochromocytoma (PC-12) cells with a low dissociation constant (kD) value of 10^-11^ M [20]. *In vitro* conditions, the peptides showed significant neuroprotective activity against PT-induced neurotoxicity in PC-12 cells [20].

Inspired by the above study’s findings [20], we investigated the neuroprotective properties, including the mechanism(s) of custom peptides [TNP and HNP] against PT-induced nerve degeneration in an *in vivo* model. The general acceptance of *Caenorhabditis elegans* as an *in vivo* model in neuronal research to comprehend neural lineage development and neuronal differentiation is noteworthy. With its transparent body, small size, short life cycle, and conspicuous and well-developed neural system, *C. elegans* is a popular model organism for neurological research because of these features [21–24]. Unlike experimental rodents, they do not require any room for growth, their maintenance is easy and cost-effective, and they have high ethical value for using laboratory experimental animals. A transgenic strain (BZ555; Pdat-1::gfp) has dopaminergic (DAergic) neurons expressing green fluorescence protein (GFP), and has been used to study neurodegeneration. Another *C. elegans* strain (NL5901; Punc-54:: α-synuclein:: YFP+unc-119) expressing the human α-synuclein protein tagged with yellow fluorescence protein (YFP) in the muscles (one of the critical proteins involved in PD), can easily be live imaged by *a* confocal microscope. Therefore, neuronal damage caused by toxic substances and their regeneration by a therapeutic molecule can be assessed speedily in *C. elegans* model [25–28]. Further, the genome of this worm is wholly sequenced and shows 60-80% similarity with human disease genes, which is an added advantage of using them as *in vivo* model organisms [29–31].

Therefore, in this study, different strains (wild type and mutant) of *C. elegans* were used to investigate the therapeutic role of custom peptides to prevent PT-induced DAergic neuron degeneration, chemosensory behavior, mitochondrial dysfunction, apoptosis, and α-synuclein protein aggregation (a hallmark for the progression of PD). RT-PCR, transcriptome, and functional proteomic analyses revealed the mechanism behind the tailored peptides’ *in vivo* neuroprotective potential in *C. elegans*. This study showed that pre-treatment with tailored peptides could block skn-1 pathway; this potential was considerably enhanced with PT treatment. In addition, a network of *C. elegans’* gene expression profiles was deciphered to comprehend the molecular relationships among the regulatory proteins and repair the harm caused by PT caused by the bespoke peptides.

## 2. Materials and Methods

### 2.1. Materials and reagents

S Biochem, Thrissur, Kerala, India synthesized custom peptides [TNP (NH2-GGDRCSGGKVGGK-COOH) and HNP (NH2-HGGKGIGKGGTGGAGVG-COOH)], a scrambled peptide (NH2-KGGDRCSGGAVGVK-COOH) and fluorescein isothiocyanate (FITC)-conjugated peptides. Mouse NGF 2.5S was obtained from Sigma Aldrich, USA. *C. elegans* Bristol wild-type strain N2, NL5901, BZ555, and CAM-1(ak37) mutant strain, the bacterial control food *Escherichia coli* (*E. coli*) OP50 (OP50), were purchased from Caenorhabditis Genetics Centre (CGC), University of Minnesota, USA. CAM-1(ak37) mutant strain was a kind gift from Dr. Kavita Babu, Associate Professor, Indian Institute of Science, Banglore. All chemicals were of analytical grade from Merck (Germany), Sigma (USA), HiMedia (India), Thermo Fisher Scientific, and Invitrogen USA.

### 2.2. Maintenance of *C. elegans* strain

Wild-type Bristol strain N2 was used as an *in vivo* model for determining PT-induced oxidative stress and its reversal by the custom peptides. The transgenic BZ555 (Pdat-1::gfp) strain of *C. elegans* has DAergic neurons expressing GFP, which was used for studying PT-induced neurodegeneration and protection by the custom peptide. A Transgenic NL5901 strain of *C. elegans* (Punc-54:: α-synuclein:: YFP+unc-119; expressing human α-synuclein protein tagged YFP in the muscles) was used to measure the deposition of toxic α-synuclein protein after custom peptides treatment. *C. elegans* (N2), BZ555, CAM-1(ak37), and NL5901 strains were grown on nematode growth medium (NGM) plates at 20 ^◦^C in a bio-incubator, and *E. coli* strain OP50 was provided as a food source. Synchronization was performed using the standard procedure of the alkaline hypochlorite method described previously [32] with some modifications. The NGM plates with adequate eggs were washed with M9 buffer and collected into a 15 mL conical centrifuge tube. A mixture of sodium hypochlorite (1 mL) and 5 N NaOH (0.5 mL) was added to the 15 mL tube containing worms in 3.5 mL M9 buffer. The tubes were then agitated for 2 min and centrifuged at 7500 rpm at room temperature to settle the eggs. The supernatant was discarded, and the pellet was washed two times with M9 buffer. Then 0.1 mL of M9 buffer was added to the pellet, transferred to the unseeded NGM plate, and incubated at 20°C after adding *E. coli* OP50 to the NGM plate.

### 2.3. Predicting the binding affinity of synthetic peptides to CAM-1 receptor in *C. elegans* by *in silico* and *in vivo* analysis

#### (A) *In silico* analysis

The custom peptides were initially developed to target the Human NTRKA. To investigate the binding of these custom peptides, scrambled peptide, and mouse NGF 2.5S, a homologous *C. elegans* protein was used. Human NTRKA protein sequence was obtained from UniProt (ID: P04629), and homologous proteins in *C. elegans* were identified using BLAST search. The structural and functional information about the homologous protein found by BLAST search was retrieved from UniProt. Three dimensional (3D) structures of different protein domains in *C. elegans*. TRK (CAM-1) were manually modeled using template-based homology modeling and Alpha Fold Prediction. Furthermore, the binding sites of these domains were predicted in PrankWeb 3 server [33]. Similarly, other proteins were selected based on the neuroprotective pathways from previous studies [20]. The proteins with available 3D structures in PDB were preferably taken. However, the unavailable 3D structures were modeled using homology modeling. The structure of mouse NGF 2.5S was acquired from PDB (ID 1BET) and prepared for a study of protein-protein interactions with a modeled CAM-1 immunoglobulin (IG) and frizzled (FZ) domains model.

The peptides were processed by molecular dynamics simulation using our previously described method [20], and different properties, such as secondary structure and beta-hairpin loop formation, were analyzed. Subsequently, all the proteins were processed for peptide docking by adding hydrogens, filling missing side chains and residues, and restraining minimization. The custom peptide/mouse NGF 2.5S/scrambled peptide docking with CAM-1 receptor was carried out in the CABS Dock server [34]. The stability of the docked complexes was further analyzed by molecular dynamic simulation. The simulation was conducted in Schrodinger Desmond (academic version v2022) with an OPLS4 force field. The simulation system was prepared with orthorhombic boundary condition with 10 Å buffer region and solvated with TIP3P water model, followed by the addition of 0.15 M sodium chloride. The system was equilibrated at 300 K and 1 bar using NVT and NPT ensembles. Nose Hoover chain thermostat and Martyna Tobias Klein barostat were used with isotropic coupling at 1 ps and 2 ps relaxation time. The system was relaxed using default options provided by Desmond, and finally, production MD was carried out for 100 ns. The binding free energy of the complexes was calculated using the MM-GBSA method.

#### (B) Determination of *in vivo* binding of FITC-conjugated custom peptides to CAM-1 mutant and N2 strains of C. *elegans*

The peptides were conjugated with FITC following the procedure earlier described by Islam et al. (2020) [14]. The binding assay was performed with the synchronized wild-type L4 larvae of N2 worms and CAM-1 mutant; 10 worms for each treatment were then collected in a 1.5 mL tube and washed multiple times with M9 buffer to remove the *E. coli* contamination. The N2 worms were then treated with FITC-conjugated peptides (TNP and HNP) for 1 to 4 h and the CAM-1 mutant worm for 2 h at room temperature, followed by washing the worms with M9 buffer (to remove adherent *E. coli*) and transferred onto 3% (w/v) agarose pads on a glass slide for confocal microscopy study (TCS SPE, Leica, Wetzlar, Germany). The excitation wavelength was set at 488 nm (Jex ∼488 nm), and the emission fluorescence signal was recorded at 519 nm (Jem∼519 nm) to detect the binding of peptides to *C. elegans*. The image was then quantified using Image J software.

The experiment mentioned above was performed to determine the binding site of custom peptides. The worms were divided into the following groups:

i. Only FITC (50 µg/mL) treated N2 worms for 2 h.
ii. Only FITC-custom peptide (HNP, 50 µg/mL or 35.7 µM) and FITC-conjugated scrambled peptide (50 µg/mL or 38.5 µM) treated N2 worms for 2 h.
iii. FITC-conjugated custom peptide (HNP, 50 µg/mL or 35.7 µM) pre-treated N2 worms for 2 h, followed by PT (10 mM) treatment for 1 h.
iv. PT (10 mM) treatment for 1 h followed by FITC-conjugated custom peptide (HNP, 50 µg/mL or 35.7 µM) treatment for 2 h in N2 worms.
v. FITC-conjugated custom peptide (HNP) co-treated N2 worms with different concentrations of PT (5 mM, 7.5 mM, 10 mM) were observed under a confocal microscope.
vi. FITC-conjugated peptides (HNP, 50 µg/mL) treated with CAM-1 mutant strain for 2 h. The signals were recorded at an emission wavelength (Jem) of ∼519 nm and quantified using Image J software.

### 2.4. Assessment of the protective role of custom peptides against PT-induced toxicity in *C. elegans* (N2 and CAM-1 mutant strain)

The *in vivo* protective activity of custom peptides (TNP, HNP, and scrambled peptide) and mouse NGF 2.5S was determined against the PT-induced oxidative stress in the *C. elegans* N2 and CAM-1 mutant strains [35]. Synchronized L4-stage adult N2 worms were pre-incubated with different concentrations (12.5 µg/mL to 100 µg/mL; equivalent to 9.6 to 76.9 µM) of custom peptides. Mouse 2.5S-NGF (50 µg/mL), vitamin C (100 µg/mL), and quercetin (50 µg/mL) were used as a positive control, whereas scrambled peptide, which did not show binding with CAM-1 receptor was used as a negative control. The N2 larvae were pre-treated with them for 2 h followed by PT (10mM)-treatment for 1 h. In an additional set of experiments, the CAM-1 mutant was pre-incubated with 50 µg/mL of custom peptides (HNP / TNP) and mouse NGF 2.5S for 2 hours, followed by PT treatment for 1 hour, to demonstrate the binding of custom peptide and mouse NGF 2.5S (at optimal dose and timing) to the CAM-1 receptor.

In another set of time-dependent studies, N2 worms were pre-incubated with custom peptides and quercetin (50 µg/mL, positive control) for 2, 12, and 24 h for 2 h (optimum pre-incubation time, see below). For 24 h of pre-incubation study, the custom peptides were added at 12 h of pre-incubation. The pre-treated worms were then transferred into the wells of 96-well plates (20 worms per well in triplicates; approximately 60 individuals for each group). The worms were provided with *E. coli* OP50 (as a food source) and treated with 10 mM PT for 24 h [36, 37], whereas, in control wells, 1X PBS was added. ‘Worms’ percentage (%) survival was scored visually using a stereo-zoom microscope after 24 h. Worms that did not show movement after light exposure and gentle tapping were considered dead.

In another set of experiments, the N2 worms were treated with 10 mM PT for 1 h, followed by post-treatment with various concentrations (12.5 µg/mL to 100 µg/mL; equivalent to 9.6 to 76.9 µM) of custom peptide (HNP / TNP) and vitamin C (100 µg/mL) /quercetin (50 µg/mL, positive control)/ mouse NGF 2.5S (50 µg/mL)/ NGM buffer with OP50 (control) for 2 h and 24 h respectively at 20 °C. The survival of N2 worms was determined, as stated above. This experiment was performed in triplicate to ensure reproducibility.

### 2.5. PT-induced loss of chemotaxis behavior in *C. elegans* and its restoration by treatment with custom peptides

The chemotaxis/memory learning assay was done as described by [38] with slight alterations. Synchronized wild-type N2 strain of *C. elegans* (L4 stage) was subjected to the following treatment conditions at 20 °C: (i) untreated (control), (ii) 10 mM PT-treatment for 1 h, (iii) PT (10 mM)-treatment for 1 h followed by treatment with custom peptides HNP and TNP (50 µg/mL [41.6 µM (TNP) and 35.7 µM (HNP)]) for 2 h/ vitamin C (100 µg/mL) for 24 h, (iv) custom peptide (HNP / TNP, 50 µg/mL) pre-treatment for 2 h/ vitamin C(100 µg/mL) pre-treatment for 24 h followed by PT (10 mM) treatment for 1 h, and (e) treatment with only custom peptides (50 µg/mL) for 2 h. The concentration and incubation time of the custom peptides used in this experiment were determined from the previous experiments (described in sections 4.3 and 4.4). The treated and untreated (control) worms were then collected in 60 mm NGM plates, and their chemotaxis behavior was determined below [38].

The plates were divided into two halves; one side was marked as “attractant” (A), and the other half was marked as “control” (C). For this experiment, 0.1% benzaldehyde dissolved in 100% ethanol was considered an odorant for the test sample, and 100% ethanol was an odorant for the control sample. On the “A” side, 2.5 μL of 0.5 M sodium azide containing 2.5 μL of odorant was added, whereas 2.5 μL of 100% ethanol containing 2.5 μL sodium azide was added on the opposite “C” side (Supplementary Fig S1). Immediately after the addition, 20 worms were transferred to the center of the plate. This experiment was performed in triplicates to ensure reproducibility. The assay plates were then scored for the number of worms in each quadrant after 1 h. The CI was calculated as follows:

Number of worms at the “A” quadrant ------------- (T)

Number of worms at the “C” quadrant ------------- (C)

Total number of worms taken on the plate ---------- (N)

Therefore, CI = (T - C)/N

### 2.6. Determination of the effect of custom peptides in inhibiting the reactive oxygen species (ROS) production in PT-treated *C. elegans*

The ROS level in *C. elegans* (N2 and CAM-1 mutant) post-PT treatment was determined using the fluorescent probe chloromethyl ‘2’,7’-dichlorofluorescein-diacetate (CM-H_2_DCFDA) by following the procedure of Kumar et al. [37]. Briefly, the N2 worms were synchronized, as stated above. Worms were washed and re-suspended in 1X PBS containing 50 µg/mL peptide (HNP / TNP). The mouse NGF 2.5S (50 µg/mL) and vitamin C (100 µg/mL) were positive controls. The worms were then shaken on a tube rotator (ROTOSPINTM, Tarson, India) for 2 h with custom peptides and 24 h with positive controls at 50 rpm, washed with 1X PBS, and further re-suspended in PBS containing PT (10 mM) /1X PBS (control) and incubated for 1 h at 23 °C. The worms were washed with PBS and re-suspended in 200 µL PBS. The worms were freeze-cracked and sonicated for 10 s. The worm lysate was then kept on ice for 30 min, centrifuged at 13,500 rpm for 30 min, and the supernatant was transferred to another tube. After protein estimation (Pierce™ BCA Protein Assay Kit), 25 μg of protein from each group was taken and made up the final volume of 50 μL with 1X PBS. For determining the PT-induced ROS generation, 50 μL protein extract containing 25 μg of each group was mixed with 100 μL of 50 μM of CM-H_2_DCFDA in PBS and incubated for 4 h at 37°C. Finally, fluorescence was measured in a multimode fluorescence plate reader (excitation at 485 nm and emission at 535 nm). A hundred worms per treatment group were used to measure ROS production. This experiment was performed in triplicates to ensure reproducibility.

The confocal microscopic analysis also quantified the *in vivo* cytoplasmic ROS level in another set of experiments. For this experiment, ten worms of each wild type and CAM-1 mutant strain were divided into three groups. They were treated as–(i) worms pre-treated with peptides (50 μg/mL) for 2 h and mouse NGF 2.5S (50 μg/mL)/ vitamin C (100 μg/mL, positive control) for 24 h followed by PT (10 mM) treatment for 1 h, and (ii) only PT (10 mM) treated worms for 1 h, and (iii) worms incubated with 1X PBS (control) for 3 h. Each group of N2 and CAM-1 mutant worms was then incubated with 50 μM of CM-H_2_DCFDA at 37°C for 4 h, visualized under a confocal microscope (TCS SPE, Leica, Wetzlar, Germany). Images were photographed at 40X with a CCD camera, and the fluorescence intensity was determined and quantitated using Image J software. The experiments were carried out in triplicates to ensure reproducibility.

### 2.7. Determination of the effect of custom peptides on reducing the PT-induced depolarization of mitochondrial membrane potential

Mitoprobe JC-1 assay kit (Invitrogen) was used to monitor mitochondrial transmembrane potential using the Mito Probe™ JC-1 Assay Kit protocol. The L4-stage young adult of each N2 and CAM-1 mutant worms were divided into three groups- (i) pre-incubated with 50 µg/mL peptides for 2 h and vitamin C for 24 h, mixed well by shaking on a tube rotator for 2 h at 50 rpm followed by the treatment with PT (10 mM) for 1 h at 20 °C, (ii) 1X PBS (control) treated worms for 3 h, (iii) only PT (10 mM) treated worms for 1h at 20 °C. After the treatment, worms were washed three times, incubated with 1 μg/mL JC-1 dye, and incubated at room temperature for 4 h. Carbonyl cyanide m-chlorophenyl hydrazone (CCCP, mitochondrial uncoupler) was used as a positive control. The experiments were repeated in triplicates to ensure reproducibility. The worms were washed three times, and the images were captured at excitation and emission wavelengths of 490 nm and 530 nm, respectively, for green fluorescence JC-1 monomers and at excitation and emission wavelengths of 525 nm and 590 nm for red fluorescence J-aggregates under a confocal microscope. The fluorescence intensity was measured using Image J software.

### 2.8. Quantitative analysis of the effect of custom peptides on PT-induced dopaminergic (DAergic) neurodegeneration

The transgenic BZ555 (P^dat-1^::gfp) strain of *C. elegans* has specifically DAergic neurons expressing GFP, to which PT could induce degeneration [39, 40]. Briefly, the L4 stage nematode of the BZ555 strain was washed with PBS and collected in a 1.5 mL centrifuge tube containing M9 buffer. The worms were treated under four different conditions- (i) 1X PBS (control), (ii) 10 mM PT-treated worms for 1 h, (iii) PT (10 mM)-treatment for 1 h followed by treatment with custom peptide (HNP / TNP, 50 µg/mL) for 2 h, and (iv) custom peptides treatment for 2 h and vitamin C (100 µg/mL) for 24 h followed by PT (10 mM) treatment for 1 h. A confocal microscope was used to view and photograph the worms, and Image J software was used to analyze the fluorescent images. The experiment was repeated three times to ensure reproducibility.

### 2.9. Quantitative analysis of the effect of custom peptides on preventing the α-synuclein aggregation in the NL5901 strain of *C. elegans*

NL5901 worms were used to observe the aggregation of the α-synuclein protein after being treated with custom peptides (50 µg/mL), as stated previously [41]. Briefly, synchronized L4 stage nematodes were incubated with 50 µg/mL custom peptides for 12 h and 100 µg/mL vitamin C for 24 h at 20 °C. After treatment, worms were washed two times with M9 buffer and transferred onto a 3 % agarose pad slide containing 100 mM sodium azide. The fluorescence intensity of the accumulated α-synuclein was observed under a microscope, and the live worms were also observed under a confocal microscope. The fluorescent and confocal images were photographed and analyzed using Image J software. The experiment was repeated three times to ensure reproducibility.

### 2.10. Quantitative reverse transcription-polymerase chain reaction (qRT-PCR) analysis to determine the effect of pre-treatment of *C. elegans* with custom peptides on the PT-induced stress-related gene expression

The procedure for qRT-PCR analysis of stress-related gene expression is adopted from our previous studies [37, 42]. Approximately 400-500 L4-stage adult worms were collected in a 1.5 mL centrifuge tube and washed multiple times to remove the adhering bacteria. Further, the worms were centrifuged, the supernatant was aspirated, and the pellet was re-suspended in 1 mL of 1X PBS containing 50 µg/mL peptide (HNP / TNP) and 100 µg/mL vitamin C (positive control), and shaken on a tube rotator for 2 h and 24 h respectively at 50 rpm. The worms (pellet) were then washed, and the pellet was re-suspended in 1 mL of 1X PBS containing 10 mM PT and incubated for 1 h at 20 °C. Then, the worms were washed, total RNA was isolated using an RNA isolation kit (Invitrogen, USA), and RNA quality was determined by measuring absorbance at 260/280 nm in a nanodrop spectrophotometer. The cDNA was described from mRNA as described previously [20]. The sequence of the primers used for qRT-PCR [37] is shown in Supplementary Table S1. The relative expression of every gene was analyzed by normalizing it with the expression of the *act-1* housekeeping gene using the 2^−ΔΔCt^ method under identical experimental conditions. The experiment was repeated three times to ensure reproducibility.

### 2.11. Transcriptomic analysis to study the gene expression in *C. elegans* when treated with PT vs. pre-treatment with custom peptides followed by treatment with PT

Synchronized L4 stage wild-type N2 strain of *C. elegans* was subjected to the following treatments at 20 °C: (a) CT group, (b) PT group, (c) PHNP, (d) HNP group. Total RNA extraction was done from the treatments of C. *elegans* using the Trizol Method [43]. Extracted RNA was subjected to DNase treatment per manufacturer protocol using New England Biolabs (NEB) (Cat# M0303L kit). The concentration and quality of RNA were measured using a Qubit fluorometer. Further, the quality was also examined by checking the RNA integrity number (RIN) using BioAnalyser from Agilent. The samples with RIN value ≥ 7 were subjected to the RNA library preparation.

Following the manufacturer’s protocol, transcriptomic mRNA libraries were prepared using NEB (cat# E7770L kit) as previously described [44]. The libraries were subjected to quantification using Qubit and Bioanalyser from Agilent. Sequencing done on an Illumina 150 bp PE platform was outsourced (Biokart India Pvt Ltd, Banglore, India).

The quality of reads obtained from the libraries was assessed using Fastqc (v0.11.8) followed by eliminating adapters / inferior-quality reads using Trim Galore (v 0.6.7). Trinity (v v2.13.2) was used for *de novo* transcriptome assembly. The redundancy of the primary assembly was removed and validated using cd-hit and RNA-Seq by Expectation-Maximization (RSEM), respectively.

The validated transcripts are subjected to Gene Ontology (GO) analysis, a Database for Annotation, Visualization, and Integrated Discovery (DAVID), and respective databases. Raw read count files were obtained by using tools viz RSEM (v v1.3.3) / Spliced Transcripts Alignment to a Reference (STAR) (v v2.7.10a) or Salmon (v v1.6.0)/ Binary Alignment Map (BAM) & Sequence Alignment Map (SAM) (v v1.14)/ Picard (v v2.18.7). For a comparative analysis, the Interactive Gene Expression Analysis Kit for Microarray & RNA-seq Data (iGEAK) (v1.0a) tool was used, which is an R (v3.3.2) and JavaScript-based open-source desktop application with the Shiny platform. Further, Voom /edgeR/limma analysis was performed to analyze differentially expressed genes (DEG). The data generation and analysis were performed at Biokart India Pvt Ltd, Bangalore, India.

### 2.12. Quantitative proteomics analysis to compare the expression of global proteins between *C. el*egans pre-treated with custom peptides followed by PT and *C. el*egans treated only with PT

For comparison of differential expression of the cellular proteins, the worms were subjected to the following treatments at room temperature: (a) CT group (untreated), (b) PT group (treated with PT), (c) PHNP group (pre-treated with HNP for 2h followed by PT treatment for 1h, and (d) HNP group (only HNP treatment. Approximately 400-500 L4-stage adult worms from each group were collected in a 1.5 mL centrifuge tube and washed multiple times to remove the adhering bacteria. Further, the worms were centrifuged, the supernatant was aspirated, and the pellet was re-suspended in 1 mL of 1X PBS. The proteins were extracted separately from the above-mentioned treated groups using RIPA lysis buffer containing proteinase K inhibitor for 30 min on ice, followed by sonication and centrifugation.

The extracted proteins were quantified, and 30 µg of the protein from each treatment group was subjected to ESI LC/MS-MS analysis as described in our previous study [43]. Briefly, the proteins are reduced (10 mM DTT) and alkylated (50 mM iodoacetamide), followed by subjected to digestion with sequencing-grade trypsin for 18 h at 37°C. The tryptic-digested peptides were desalted, concentrated, and subject to nano-UHPLC, followed by LC/MS-MS analysis. For ionization and fragmentation nano spray electrospray positive ionization (ESI) and collision-induced dissociation were used, respectively. MS and MS/MS scans (ranging from 500 to 2000 m/z) were carried out in FT-ICR/Orbitrap and linear ion trap, respectively. Only double or triple-charged ions were selected for collision-induced dissociation (CID) MS/MS analysis.

The raw data obtained from MS/MS analysis was analyzed in Peak studio, and the parameter for identifying proteins was adopted from our previous study[20]. The relative abundances were calculated from the spectral intensity of the respective protein, and fold change values of the relative abundances were also calculated.

### 2.13. Determination of acute *in vivo* toxicity and biochemical changes post-treatment of the peptides in a Swiss albino mouse model

Acute *in vivo* toxicity of custom peptides was assessed in Swiss albino mice (18-20 g) following OECD guidelines. Ethical permission was approved by the animal ethics committee of IASST (IASST/IAEC/2022/10). For *in vivo* toxicity evaluation, custom peptides (TNP: HNP, 1:1 w/w) were dissolved in 0.2 mL of 1X PBS (pH-7.4) and injected intravenously (10 mg/kg body weight, *i.v.*) into the Swiss albino mice (n=6) (70). The control mouse group received only 0.2 mL of 1X PBS (pH-7.4) (placebo). The treated mice were continuously monitored at intervals of 6 h up to 24 h of injection for any behavioral alteration or death. After the mice were treated with customized peptides for 24 h, they were sacrificed, and blood was extracted immediately via heart puncture. Using a semi-automatic biochemistry analyzer, the BeneSphera C61, plasma was separated from the blood (control/treated group) by centrifuging it at 4500 rpm for 20 min at 4°C. The biochemical parameters that were measured included serum glutamic oxaloacetic transaminase (SGOT), alkaline phosphatase (ALKP), serum glutamic pyruvic transaminase (SGPT), BUN (blood urea nitrogen), glucose content, creatinine, cholesterol, bilirubin, and albumin level.

To study possible custom peptides (1:1)-induced morphological changes, the heart, liver, brain, kidney, liver, ovary, and testis of each control and treated group of mice were dissected after 24 h of treatment. Tissues were fixed in 10% buffered formaldehyde, dehydrated, and embedded in paraffin. The slides were observed under light microscopic (Zeiss Axiolab A1) after hematoxylin-eosin (H/E) staining (70). Serum levels of proinflammatory cytokines (TNF-α, IL-6, IL-1β) were determined by a commercial immunoassay kit (Quantikine® HS Immunoassay kit, biotech R and D systems) and followed manufacturer instructions.

### 2.14. Statistical analysis

The data are presented as mean ± standard deviation (SD) of triplicates. Student’s t-test (Sigma Plot 11.0 for Windows 10.0) was to compare between the test and control and for more than two groups analysis of variance (ANOVA) was performed followed by post hoc analysis (GraphPad Prism software). The *p*-value ≤ 0.05 was considered statistically significant.

## 3. Results

### 3.1. *In silico* analysis of custom peptides binding in *C .elegans*

According to the modelling results, mouse NGF 2.5S and custom peptides (TNP and HNP) bind to a CAM-1 receptor domain that resembles the mammalian TrkA receptor and is immunoglobulin (IG)-like. Despite the substantial variations in the total amino acid sequences, the domains of both the proteins share a comparable structural similarity asthe amino acid sequences of the binding regions of both the proteins are similar. The tyrosine-protein kinase receptor CAM-1’s IG domain was found to interact with the custom peptides TNP (Fig. 1a), HNP (Fig. 1b), and mouse NGF 2.5S (Fig. 1c). Nevertheless, the IG domain of the CAM-1 receptor did not show appreciable interaction with scrambled peptide (Fig 1d).

**Fig 1.**
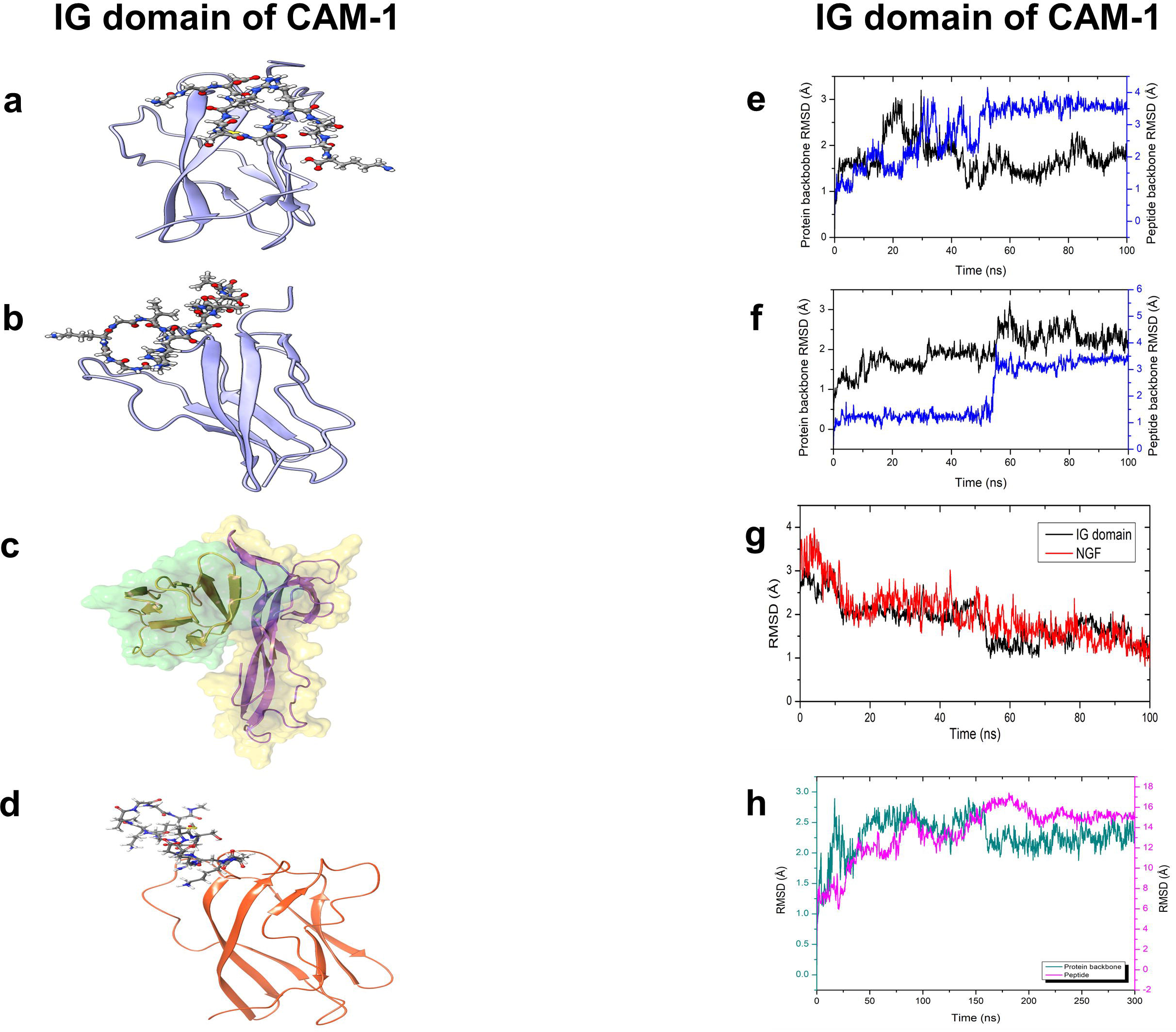
Custom peptide (a) TNP, (b) HNP, (c) mouse NGF 2.5S, and (d) scrambled peptide complex with CAM1-IG domain shown in cartoon representation. Root mean square deviation (RMSD) plot of (e) TNP, (f) HNP, and (g) mouse NGF 2.5S, and (h) scrambled peptide complex with IG domain.

The RMSD analysis of the trajectory demonstrated significant stable interaction for all these proteins (Figs 1e-h). The MM-GBSA binding free energies obtained for different complexes were between -40 and -51 kcal/mol. The root mean square deviation (RMSD) values exhibited variation up to 4.2 Å.

The custom peptide showed efficient binding with one of the tyrosine-protein kinase receptors CAM-1. Entries in the Reactome pathway browser database [45] suggest that CAM-1 is an essential component in the MAPK signaling pathway by playing multiple roles. Custom peptides TNP (Figs 1a and e), HNP (Figs 1b and 5.1 f), and mouse NGF 2.5S (Figs 1c and g) form a complex with IG of CAM-1 and exhibited reasonable binding stability. In contrast, scrambled peptide (Figs 1d and h) showed relatively unstable binding. Notably, BLAST sequence search revealed that CAM-1 shares considerable homology (38.5%) with human NTRKA receptors and functions similarly.

### 3.2. The confocal microscopic study shows *in vivo* binding of FITC–conjugated custom peptides to the nerve ring region of *C. elegans* N2 strain; however, an insignificant binding was observed with CAM-1 mutant strain of *C. elegans*

The confocal microscopic images of the dose-dependent (12.5 µg/mL to 100 µg/mL) and time-dependent (1 h - 6 h) binding of the FITC-custom peptides (TNP and HNP) to *C. elegans* (L4 of N2 strain) are shown in Supplementary Figs S2 a-b and Figs 2a-b respectively. The FITC-custom peptides (TNP and HNP, 50 µg/mL) showed binding at 2 h of incubation at the nerve ring part and hypodermal syncytia covering the lips and head portion of the *C. elegans* (Figs 2a-b). However, the FITC-scrambled peptide (50 µg/mL, 38.5 µM) did not show binding (2 h incubation) to the CAM-1 receptor at the nerve ring region of N2 worms (Supplementary Figs S3 a-b).

**Fig 2.**
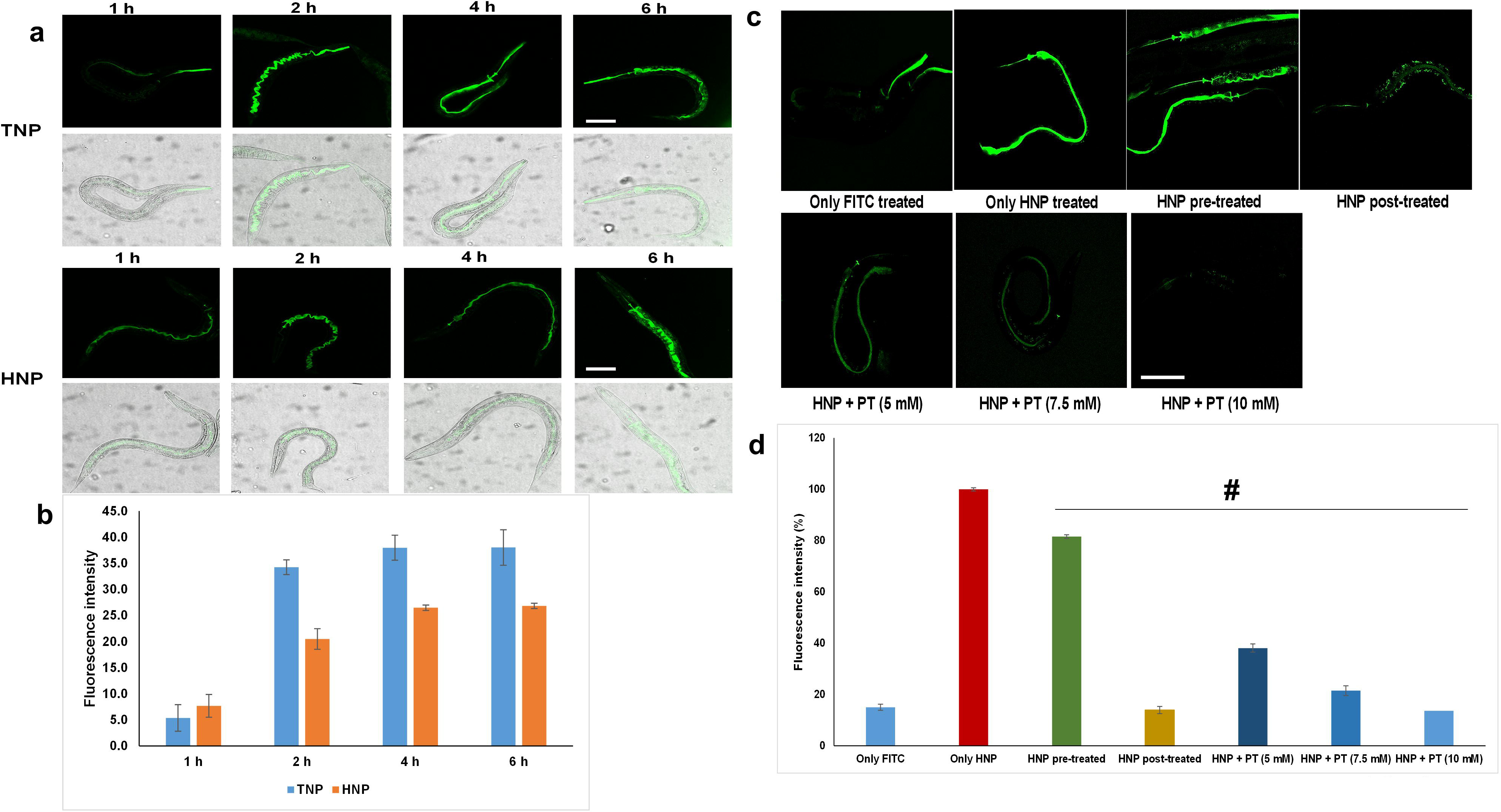
Confocal microscopic (40 X) studies of the *in-vivo* binding of FITC-custom peptides to *C. elegans*. **(a)** Time-dependent (1 h – 4 h) binding of custom peptides (50 µg/mL) to the *C. elegans*. **(b)** Bar graph representing fluorescence intensity between the treatment groups **(c)** Microscopic image of the custom peptide binding to *C. elegans* in pre-treatment, post-treatment, and co-treatment conditions (with various concentrations of PT). The scale bar indicates the length as 100 µm**. (d)** Bar graph representing fluorescence intensity between the treatment groups; Significant difference between the pre-treatment, post-treatment, and cotreatment with FITC-custom peptide HNP compared to binding of only FITC-conjugated HNP to *C.elegans*, ^#^p ≤ 0.05. Values are mean ± SD of triplicate determinations.

Pre-treatment of *C. elegans* with FITC-conjugated TNP and HNP showed optimum fluorescence intensity at the nerve ring region; nevertheless, only FITC-treatment (control) did not show fluorescence signal at the nerve ring region of *C. elegans,* indicating binding of peptides to the nerve ring region of *C. elegans* (Figs 2c-d). However, the post-addition of peptides after PT treatment showed a significant decrease in fluorescence intensity at the nerve ring region compared to custom peptides pre-treated worms before adding PT (Figs 2c-d). Further, when a fixed concentration of peptides was co-treated with increasing concentrations of PT, a PT concentration-dependent reduction in the fluorescence intensity at nerve ring regions was observed, suggesting custom peptides and mouse NGF 2.5Sprotect the nerve ring region of the *C. elegans* from PT-induced neuronal damage (Figs 2c-d).

Further, the binding efficiency of FITC-custom peptides (TNP and HNP) at their optimum dose (50 µg/mL) and time (2 h) in the N2 strain and CAM-1 mutant strain of *C. elegans* were compared in (Figs 3a-b). However, negligible binding was observed at the nerve ring region in the CAM-1 mutant strain of *C. elegnas*, compared to the binding of FITC-custom peptides in the N2 strain of *C. elegans* which is evident by a significant (*p* ≤ 0.05) decrease in the fluorescence intensity (19-27 %) of FITC-custom peptides was observed in the CAM-1 mutant strain when compared to the N2 strain of *C. elegans* (Fig. 3b).

**Fig 3.**
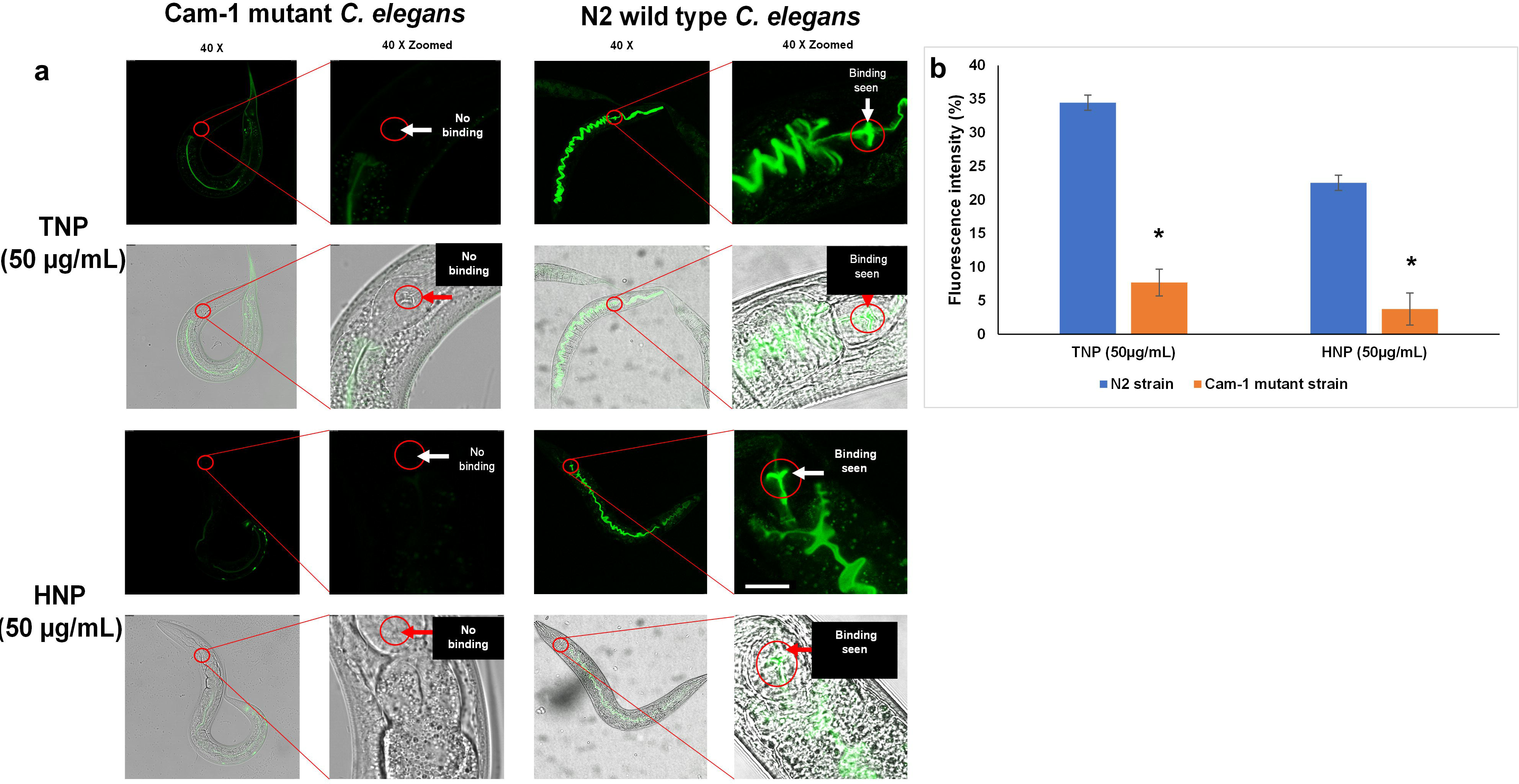
(a) Confocal microscopic (40 X) studies of the *in-vivo* binding of FITC-custom peptides (50 µg/mL, 2 h) to CAM-1 mutant and compared with wild-type N2 strain *C. elegans*. The scale bar indicates the length as 100 µm**. (b)** Bar graph representing fluorescence intensity between the CAM-1 mutant and N2 strains. *p ≤ 0.05, a significant difference between CAM-1 mutant and N2 strain of *C. elegans*.

### 3.3. Pre-treatment with custom peptides reduced PT-induced death and restored the chemotaxis dysfunction

Custom peptides pre-treatment demonstrated a concentration-dependent increase in the survivability of the N2 strain of *C*. *elegans* against PT (LC_50_ value)-induced death, and the optimum concentration was determined for both the peptides at 50 µg/mL [41.6 µM (TNP) and 35.7 µM (HNP)] (Fig 4a). HNP showed a significant (*p* ≤ 0.05) increase in survivability compared to TNP. Nevertheless, a wealth of data suggests that enhanced biological activity might not always follow from effective binding with the receptor [20, 46]. The two peptides may exhibit varied inhibition against PT-induced toxicity due to their differing binding affinities and efficacies. The non-specific or unstable binding of peptides to the receptor may cause the binding efficiency and affinity variation. The pre-treated group that received 100 µg/mL of antioxidant vitamin C (positive control) exhibited a negligible protective effect against death (Fig 4a). Neuroprotective efficacy against PT-induced toxicity in *C. elegans* was not demonstrated by the scrambled peptide pre-treatment (Supplementary Figs S4).

**Fig 4.**
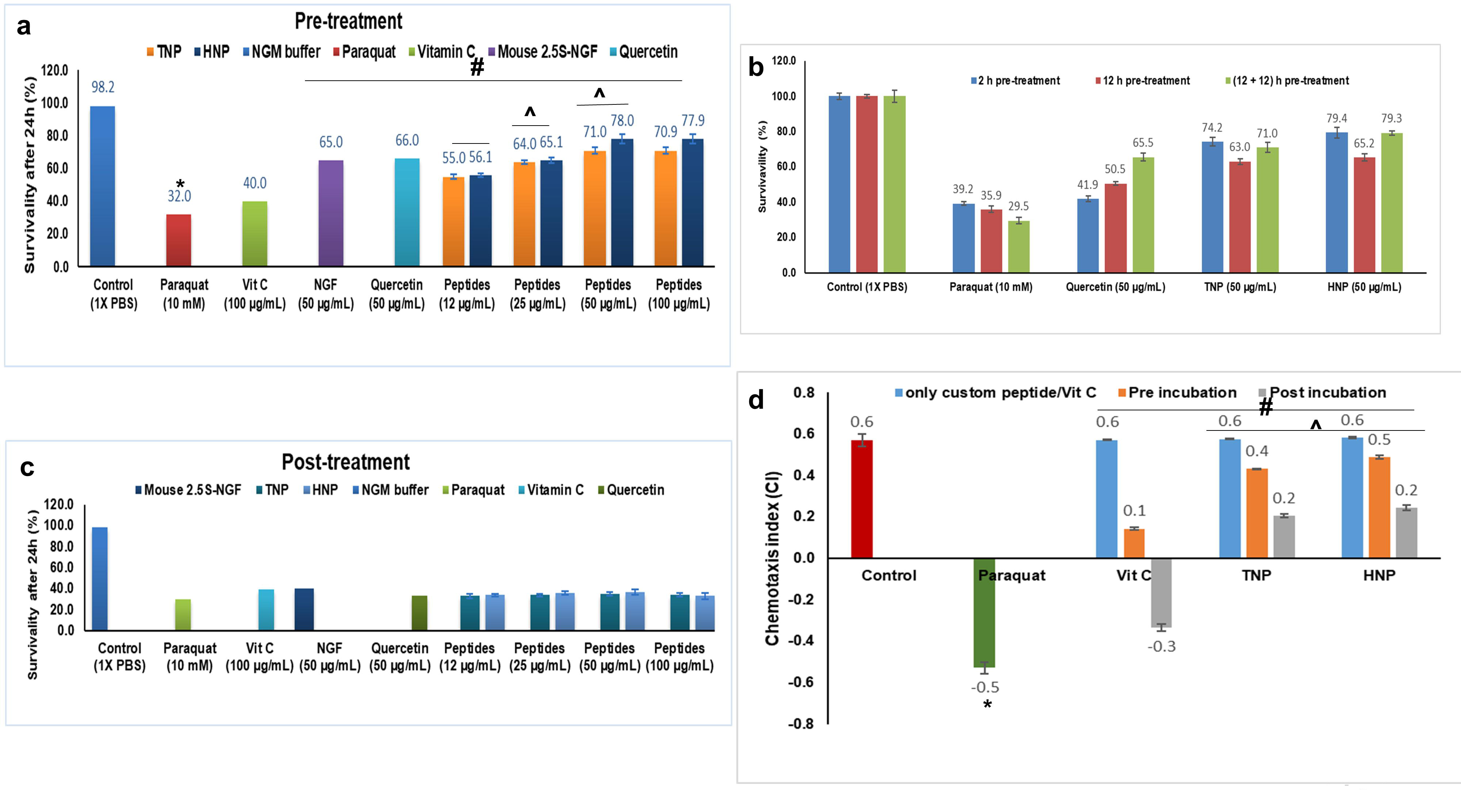
Determination of the effect of the custom peptide on PT-induced death of *C.elegans*. **(a)** worms were pre-incubated with mouse NGF 2.5S (50 µg/mL)/quercetin (50 µg/mL, positive control) / vitamin C (100 µg/mL, positive control) and progressive concentration of custom peptides (12 µg/mL - 100 µg/mL) followed by the PT (10 mM) treatment. *p ≤ 0.05, a significant difference between untreated (control) and PT-treated cells; ^#^p ≤ 0.05, a significant difference between PT-treated cells and quercetin/ mouse NGF 2.5S/ vitamin C and custom peptide pre-treated *C. elegans*. ^p ≤ 0.05 Significance of difference in different concentrations for custom peptides. **(b)** worms were pre-incubated with quercetin (50 µg/mL, positive control) and custom peptides (50 µg/mL) for 2 h, 12 h, and 24 h followed by the PT (10 mM) treatment. *p ≤ 0.05, a significant difference between untreated (control) and PT-treated cells; ^#^p ≤ 0.05, a significant difference between PT-treated cells and quercetin (positive control) and custom peptide pre-treated *C. elegans*. ^p ≤ 0.05, a significant difference between quercetin pre-treated *C. elegans* and the peptide (TNP and HNP) pre-treated *C. elegans*. **(c)** Worms were incubated with PT (10 mM) for 1 h followed by the treatment with custom peptides (12 µg/mL to 100 µg/mL). Freshly prepared custom peptides were added after 12 h of pre-incubation for 24 h pre-incubation condition. Worms were counted under a stereo zoom microscope for 30 s up to 24 h of treatments. Values are mean ± SD of triplicate determinations. **(d)** Restoration of chemosensory behavior in *C. elegans* pre-treated with custom peptides. Synchronized L4 stage *C. elegans* wide-type strain N2 was incubated with or without 50 µg/mL custom peptides. They were then subjected to PT-treatment (10 mM) and a chemosensory assay. *p ≤ 0.05, a significant difference between untreated (control) and PT-treated cells; ^#^p ≤ 0.05, a significant difference between PT-treated cells and quercetin (positive control) and custom peptide pre-treated *C. elegans*. ^p ≤ 0.05, a significant difference between quercetin pre-treated *C. elegans* and the peptide (TNP and HNP) pre-treated *C. elegans*. Values are means□±□cSD of triplicate determinations.

Custom peptide pre-treatment for two hours showed a significant (p ≤ 0.05) increase (35-40%) in the survivability of N2 strain of *C. elegans* compared to PT-treated *C*. *elegans*; however, 12 h pre-treatment compared to 2 h pre-treatment was found to be less effective in protecting the *C*. *elegans* against PT-induced toxicity and death. Nevertheless, 12 h pre-treatment with custom peptides followed by re-supplementation of custom peptides again for 12 h (total pre-incubation time 24 h) restored the survivability of *C*. *elegans* against PT, and this result was found to be similar as shown by pre-incubation with custom peptides for 2 h before addition of PT (Fig 4b). Pre-treatment with 50Jµg/mL quercetin, a known neuroprotectant compound, demonstrated much lower protection than custom peptides under similar experimental conditions (Fig 4b). Post-treatment with custom peptides, vitamin C (antioxidant), mouse 2.5 S NGF, and quercetin (after PT addition) did not reverse the PT-induced toxicity in *C*. *elegans* (Fig 4c). The binding of the peptides to the CAM-1 receptor was confirmed by the considerable (p < 0.05) decrease in the neutralization (23-27%) of PT-induced toxicity by the custom peptides and mouse NGF 2.5S pre-treated CAM-1 mutant strain compared to the N2 strain of *C. elegans* (Supplementary Fig S5).

No significant difference (p ≧ 0.05) in the chemotaxis index (CI) between the peptide-treated and untreated (control) *C. elegans* was observed (Fig. 4d). However, sensory neuron dysfunction was observed in PT-treated wild-type *C. elegans* (Fig. 4d). Notably, a significant restoration (p ≤ 0.05) of PT-induced sensory dysfunction was observed in pre- (CI = 0.43 ± 0.028 for TNP, and CI = 0.48 ± 0.024 for HNP) and post-peptide-treated worms (CI = 0.20 ± 0.014 for TNP, and CI = 0.24 ± 0.012 for TNP) compared to the worms treated with PT (Fig. 4d). Further, pre- or post-treatment with vitamin C (antioxidant) is ineffective in improving PT-induced sensory neuron dysfunction (Fig 4d). Peptides TNP and HNP demonstrated similar potency (p ≧ 0.5) in the restoration of PT-induced sensory neuron dysfunction in *C. elegans* (Fig 4d).

### 3.4. Custom peptides inhibit PT-induced ROS production and depolarization of mitochondrial membrane potential in *C. elegans* N2 strain but could not protect CAM-1 mutant strain

The spectrofluorometric determination of fluorescence intensity of 2,7-dichlorofluorescein (DCF) confirmed a significant (p ≤ 0.05) increase in the ROS level in PT (10 mM)-treated wild-type worms by 44% compared to control (untreated) worms. A significant reduction (p ≤ 0.05) in ROS level was observed in wild-type worms pre-treated with HNP and TNP (68-70%) and vitamin C (35%), mouse NGF 2.5S (33%) as compared to the PT-treated wild type worms (Fig 5a). The custom peptide HNP at a concentration of 50 µg/mL [41.6 µM (TNP) and 35.7 µM (HNP)] showed superior reduction (32-35%) (*p* ≤ 0.05) in the ROS generation, compared to TNP and vitamin C (positive control) (Fig 5a).

**Fig 5.**
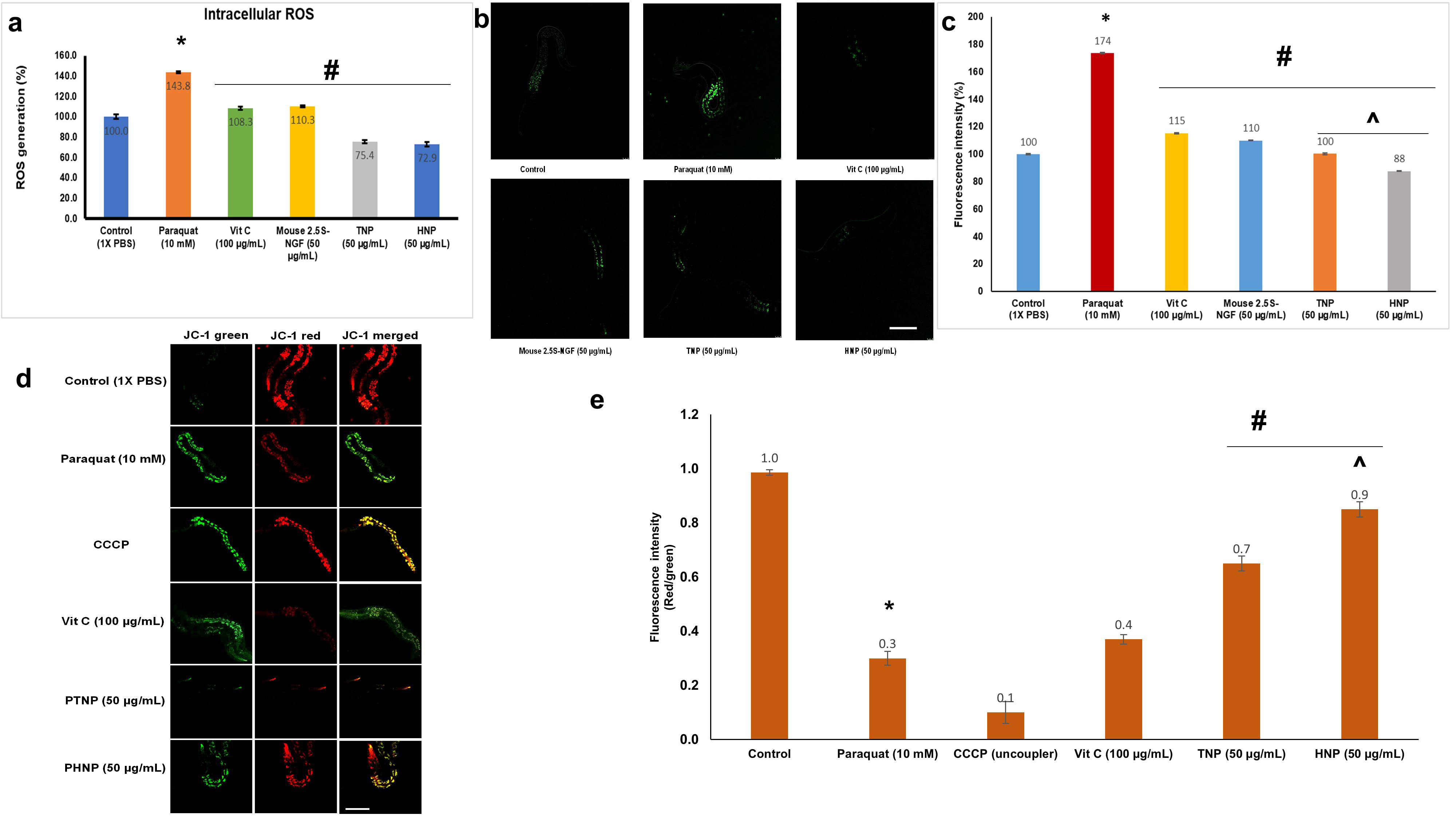
Determination of PT-induced intracellular ROS generation and its reversal by pre-treatment with custom peptide in wild type N2 strains of *C. elegans.* The ROS generation was determined by using an H_2_DCFDA fluorescence probe. **(a)** spectrofluorometric determination of intracellular ROS. *p < 0.05, the significant difference between untreated (control) and PT-treated worms; ^#^p < 0.05, the significant difference between PT-treated and custom peptide-treated worms. **(b)** confocal microscope images of nematodes expressing ROS. The scale bar indicates the length as 100 µm. **(c)** Bar graph representing dosimetry analysis of confocal images to quantitate the intracellular ROS generation. Error bars indicating SD (n=3). *p < 0.05, the significant difference between untreated (control) and PT-treated worms; ^#^p < 0.05, the significant difference between PT-treated and custom peptide-treated worms. ^p ≤ 0.05, a significant difference between Vit C/ mouse 2.5 S-NGF pre-treated *C. elegans* and the peptides (TNP and HNP) pre-treated *C. elegans*. Values are means□±□SD of triplicate determinations. **(d)** Confocal images of *C. elegans* showing reversal of PT-induced disruption of mitochondrial membrane potential (MMP) of *C. elegans* pre-treated with custom peptides. The scale bar indicates the length as 100 µm. The scale bar indicates the size as 100 µm. Carbonyl cyanide m-chlorophenyl hydrazone (CCCP) is a mitochondrial uncoupling agent that depolarises the mitochondria taken as a positive control. (The PT-treated (10 mM) N2 wild-type strain of nematodes pre-treated with or without custom peptide (50 µg/mL) was observed to measure the red/green fluorescence intensity ratio by JC-1 staining. **(e)** Bar diagram representing the red/green fluorescence intensity ratio quantified using Image J software. *p ≤ 0.05, a significant difference between untreated (control) and PT-treated worms; ^#^p ≤ 0.05, a significant difference between PT-treated and custom peptide pre-treated worms. ^p ≤ 0.05, a significant difference between the two peptides (TNP and HNP) pre-treated *C. elegans*. Values are means□±□SD of triplicate determinations.

The confocal microscopic analysis also determined that the custom peptides mediated reduction in intracellular ROS generation in wild-type N2 strain *C. elegans* (Figs 5b-c). The data showed a significant increase (p ≤ 0.05) in intracellular ROS levels in PT-treated worms compared to control wild-type worms. Further, pre-treatment of worms with custom peptides (TNP and HNP, 50 µg/mL) and vitamin C (positive control, 100 µg/mL) /mouse NGF 2.5S (50 µg/mL) significantly diminished the ROS production (Fig 5c); however, to a different extent. Custom peptide HNP, compared to TNP, showed superior activity in inhibiting PT-induced ROS production in wild-type worms (Fig 5c). No significant (p ≧ 0.05) decrease in ROS production was observed in the custom peptides pre-treatment group compared to the PT-treated group in the CAM-1 mutant strain of *C. elegans* (Supplementary Figs S6 a-b).

The JC-1 staining procedure was used to study whether custom peptides can restore the loss of mitochondrial membrane potential (MMP) triggered by PT in *C. elegans* N2 and CAM-1 mutant strains. The effect of custom peptides on MMP using JC-1 dye (cyanin dye) (Fig 5d) was demonstrated. The red/green fluorescence ratio was quantified from the confocal images to monitor the MMP (Fig 5e). The fluorescence intensity of the red/green ratio was significantly reduced (p ≤ 0.05) by 80% in 10 mM PT-treated worms compared to control (1X PBS-treated) worms (Fig 5e). However, the fluorescence intensity of the red/green ratio was significantly (p ≤ 0.05) restored by 60 to 70% when worms were pre-treated with custom peptide (TNP / HNP) as compared to PT-treated worms (Figs 5d-e), indicating a reduction of PT-induced depolarization of MMP by custom peptides in *C. elegans*. HNP showed a significant (p ≤ 0.05) 20% increase in the restoration of PT-induced mitochondrial depolarization as compared to TNP (Figs 5d-e). In contrast, vitamin C pre-treatment showed no significant difference in the restoration of PT-induced mitochondrial depolarisation (Figs 5d-e). Moreover, custom peptide pre-treatment did not offer any considerable repair of alteration of MMP in the CAM-1 mutant strain of *C. elegans* (Supplementary Figs S7 a-b).

### 3.5. Custom peptides restored PT-induced dopaminergic (DAergic) neurodegeneration and reduced α-synuclein aggregation in *C. elegans*

*C. elegans* BZ555 strain expresses a green fluorescent protein in all six intact DAergic neurons comprising two distal cephalic (CEPD), two ventral cephalic (CEPV), and two anterior deirid (ADE) (Fig 6a). Studies have shown that specific DAergic neuron degeneration is attained with 10 mM PT exposure [39, 40, 47], which showed impairment of CEPs and ADEs cell bodies (Fig 6b). The analysis showed a significant decrease (p ≤ 0.05) in the fluorescence intensity with PT treatment to 54-58% for CEPs and 70-75% for ADEs neurons compared with the PBS-treated (control) worms (Fig 6b). The TNP and HNP peptides pre-treated groups showed a significant (p ≤ 0.05) increase in fluorescence intensity for CEPs neurons and ADEs neurons (Figs 6a-b). This data suggests a considerable impairment of PT-induced neuronal degeneration and significantly higher activity of HNP as compared to TNP (Figs 6a-b). However, the vitamin C pre-treated group does not show a significant (p > 0.05) increase in fluorescence intensity for CEPs and ADEs neurons compared with the PT-treated worms (Figs 6a-b). Similarly, peptides TNP and HNP in post-treated worm groups showed a significant (p ≤ 0.05) increase in GFP intensity by 57.5-59.3% and 60.3-63.7%, respectively, for CEPs neurons and 38.5% and 65%, for ADES neurons (Figs 6a-b). Fluorescence intensity statistics were used to analyze alterations in dopaminergic neurons after CP treatment. Furthermore, the evidence indicates that ADE neurons are somewhat more successfully restored by post-treatment than CEP neurons. Consequently, there is some effectiveness during pre-treatment and following it.

**Fig 6.**
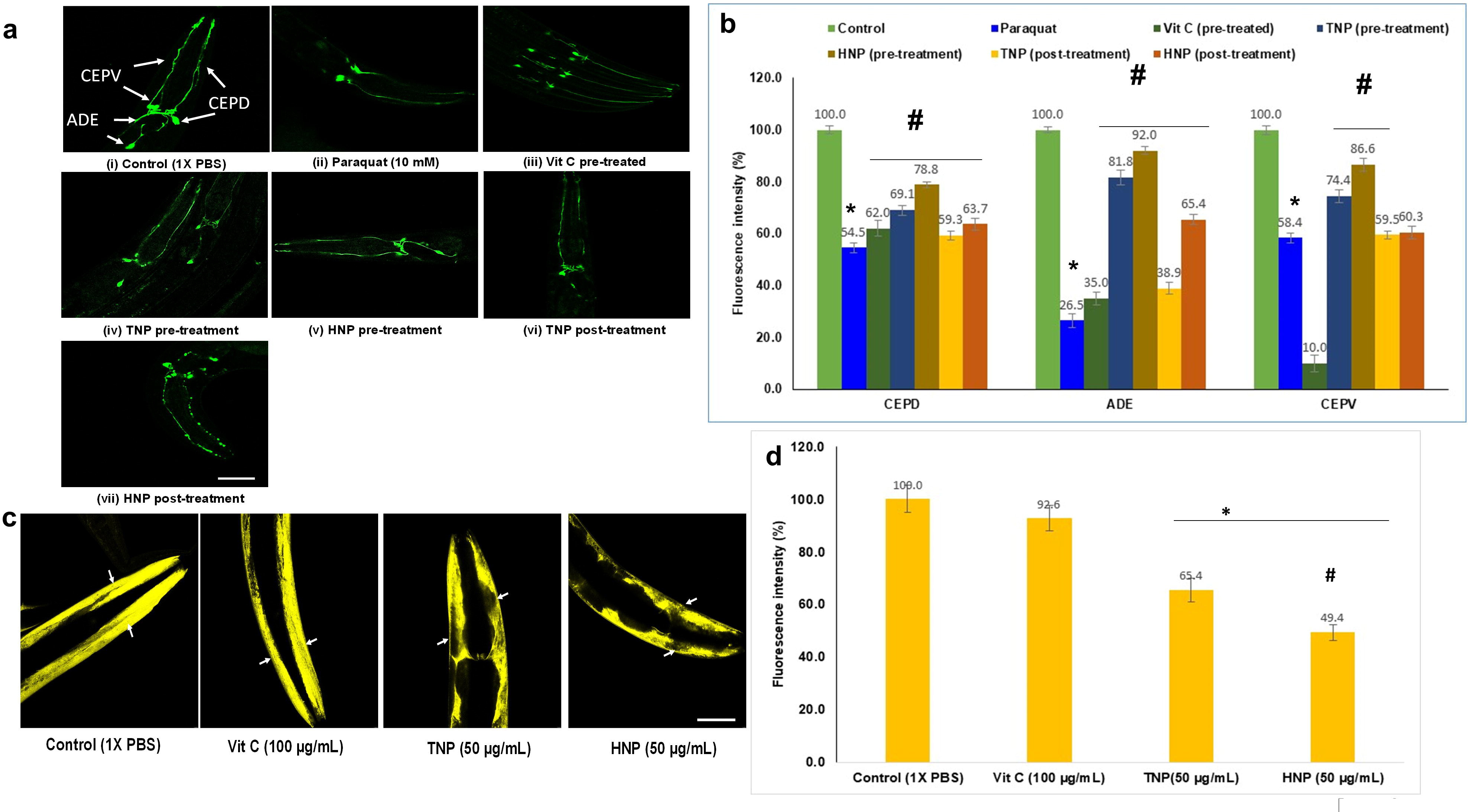
Determination of dopaminergic neurodegeneration induced by PT in BZ555 *C. elegans*. **(a)** Confocal microscopic images (40 X) of dopamine (DA) neurons emerging GFP fluorescence signals in PT-treated BZ555 worms with or without pre-treated with custom peptides (50 µg/mL). The scale bar indicates the length as 100 µm. **(b)** The bar diagram shows the GFP fluorescence intensity indicating the content of DA neurons in BZ555 worms, quantified using the Image J software. *p ≤ 0.05, a significant difference between untreated (control) and PT-treated worms; ^#^p ≤ 0.05, a significant difference between PT-treated and custom peptide-treated worms. Custom peptides inhibit the aggregation of α-synuclein in transgenic NL5901 strains of *C. elegans*. **(c)** Confocal images of custom peptides (50Jμg/mL)-treated NL5901 worms after 12 h of incubation. The scale bar indicates the length as 100 µm. **(d)** The bar chart shows the fluorescence intensity representing the α-synuclein protein aggregation in custom peptide-treated NL5901 worms for 12 h. * (p ≤ 0.05) a significant difference between control (CT) and custom peptides-treated *C. elegans*, ^#^ (p ≤ 0.05) a significant difference between TNP and HNP treated *C. elegans*. Values are means□±□SD of triplicate determinations.

The fluorescence intensity at the anterior part of the NL5901 strain of *C. elegans* showed aggregation of human α-synuclein (YFP tagged) in the body wall muscles (Fig 6c). The image analysis showed a significant decrease (p ≤ 0.05) in the fluorescence intensity in the custom peptides-treated *C. elegans* as compared with the PBS-treated (control) *C. elegans* (Fig 6d). The TNP and HNP treatment showed a significant (p ≤ 0.05) decrease in the fluorescence intensity by 37% and 60%, respectively, compared to the fluorescence intensity of the control group of worms (Figs 6c-d). HNP treatment, compared to the TNP treatment, showed a significant (p ≤ 0.05) 20% decrease in the α-synuclein aggregation in *C. elegans* (Figs 6c-d). However, vitamin C pre-treatment did not show a significant (p > 0.05) reduction in the α-synuclein aggregation, as compared to the PBS-treated (control) group of *C. elegans* (Figs 6c-d).

### 3.6. The custom peptides pre-treatment restores the PT-induced upregulated antioxidant/heat shock response/ p38 mitogen-activated protein kinase (MAPK) genes and delayed PT-induced programmed cell death in *C. elegans*

The qRT-PCR analysis data demonstrated a significant increase (p ≤ 0.05) in the expression of antioxidant genes (*sod-1*, *sod-3*, *cat-1*, *cat-3*, *trx-1*, *gst-4*, and *gst-6)*, heat shock gene *hsp-16.1*, and MAPK signaling pathways genes (*skn-1*, *sek-1*, and *pmk-1)* in PT-treated N2 worms compared to untreated worms (control). However, pre-treatment of worms with vitamin C (positive control) / custom peptides (HNP and TNP) resulted in a significant downregulation (p ≤ 0.05) or restoration of the expression of those genes upregulated by PT treatment (Fig 7). Further, the qRT-PCR analysis also showed significant (p ≤ 0.05) upregulation of pro-apoptotic genes (*ced-3* and *ced-4*) and significant downregulation (p ≤ 0.05) of antiapoptotic genes (*ced-9*) in PT-treated *C. elegans* compared to untreated (control) *C. elegans*. Further, the pre-treatment of custom peptides (TNP and HNP) restores the upregulated (*ced-3* and *ced-4*) and downregulated (*ced-9*) pro-apoptotic and antiapoptotic genes, respectively, in PT-treated *C. elegans* (Fig 7). However, vitamin C (positive control) pre-treatment did not show significant down-regulation (p ≤ 0.05) of MAPK signaling pathway genes (*skn-1*, *sek-1*, and *pmk-1)* and pro-apoptotic genes (*ced-3* and *ced-4*) compared to PT-treated worms (Fig 7).

**Fig 7.**
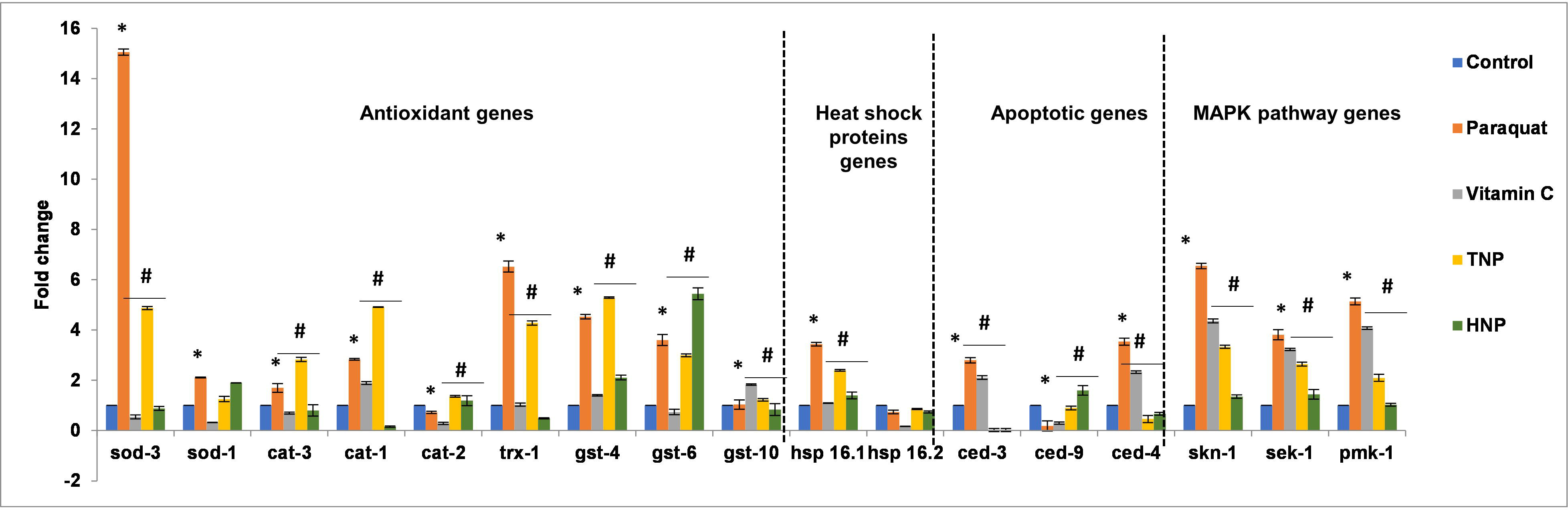
The qRT-PCR analysis shows the expression of genes involved in stress resistance, innate immunity, and apoptotic pathways in the PT-treated *C. elegans*, compared with the vitamin C (positive control)/custom peptide pre-treated *C. elegans*. The expression of mRNA was normalized using the housekeeping gene act-1. * (p ≤ 0.05) a significant difference between control (CT) and PT-treated *C. elegans*, ^#^ (p ≤ 0.05) a significant difference between PT-treated and vitamin C/ custom peptide pre-treated worms. Values are means□±□SD of triplicate determinations.

Both bespoke peptides have the same binding sites on the CAM-1 receptor, but their basic sequence and molecular weight differ slightly. Moreover, the venoms of two distinct snake species—the Indian cobra and the Russell’s viper—were used to construct them. This result suggests that the two bespoke peptides may have distinct mechanisms of action and efficacy, corroborating the research findings. This result is an exciting discovery because both peptides eventually demonstrated the ability to act as antioxidants and neuroprotectors, either by increasing the fold change of stress resistance against PT toxicity or by improving the fold change of antioxidant enzymes to remove free radicals in response to PT toxicity (Fig 7). Furthermore, qRT-PCR research revealed that HNP had higher activity than TNP. For this reason, HNP was chosen for the transcriptome and proteome studies to identify the gene regulatory networks that inhibit the neuronal deterioration caused by PT.

### 3.7. A comparison of the differential expression of mRNA between PT-treated *C. elegans* and pre-treatment of *C. elegans* with custom peptides followed by PT-treatment

The cDNA libraries were prepared for the mRNA, isolated from each group of *C. elegans* (N2) treated with (a) 1X PBS (control) treated worms (CT group), (b) 10 mM PT treatment for 1 h (PT group), (c) pre-treatment with 50 µg/mL (35.7 µM HNP) custom peptide HNP for 2 h followed by 10 mM PT treatment for 1 h (PHNP group), and (d) treatment with 50 µg/mL of custom peptide HNP for 2 h (HNP group). The mRNAs were sequenced with the Illumina 150 bp PE platform. The average total number of reads aligned was found to be 16.7 million (CT), 15.3 million (HNP), 12.8 million (PHNP), and 20.2 million (PT) for the three corresponding cDNA libraries.

The transcriptomic analysis showed differential expression of 15,102 genes in *C. elegans* compared to the PT and CT group of worms and 701 genes when compared between the PT and PHNP group, in which 441 and 260 genes were found to be upregulated and downregulated, respectively.

The transcriptomic data’s principal component analysis (PCA) revealed variance in the gene expression level between all four treatment groups (Supplementary Fig S8a). The correlation plot (Supplementary Fig S8b) shows clustering among different treated groups. Red indicates the positive, and blue indicates the negative correlation plot between the four treatment groups (Supplementary Fig S8b). The volcano (p-values vs. fold change) and scatter plots display the (i) differential gene expression between PT and CT groups (Figs 8a-b) and (ii) differentially altered genes between PHNP and PT groups (Figs 8c-d). The Volcano plot and scatter plot analyses revealed significant downregulation (Fc < -1) of most of the genes in *C. elegans* in the PT group compared to the CT group. The pre-treatment of peptide (HNP) resulted in the upregulation of those ‘genes’ expression downregulated in the PT group of worms (Figs 8a-d). The heat map analysis displayed a disparity in gene expression between the PT and PHNP groups compared to the CT group (Fig 8e).

**Fig 8.**
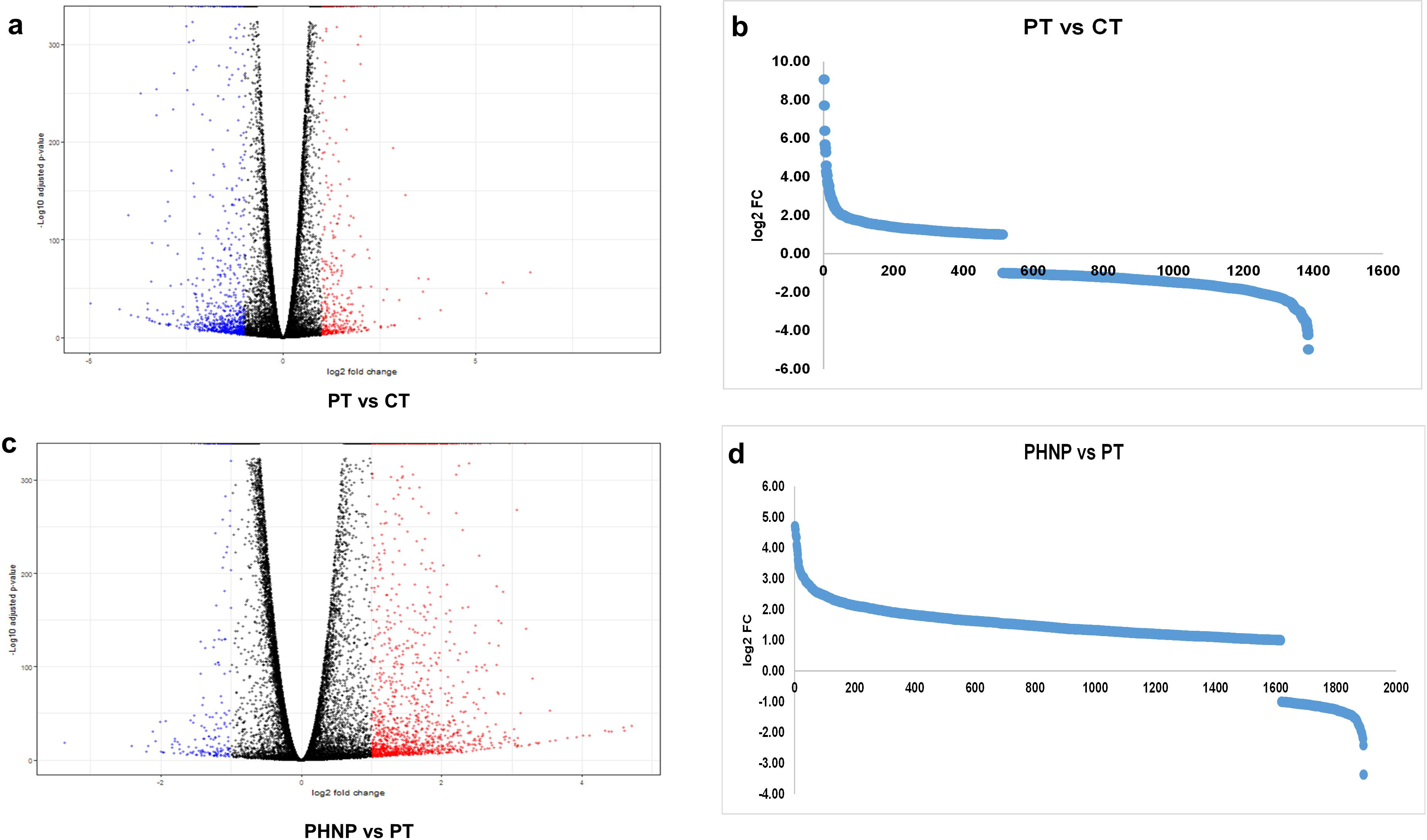
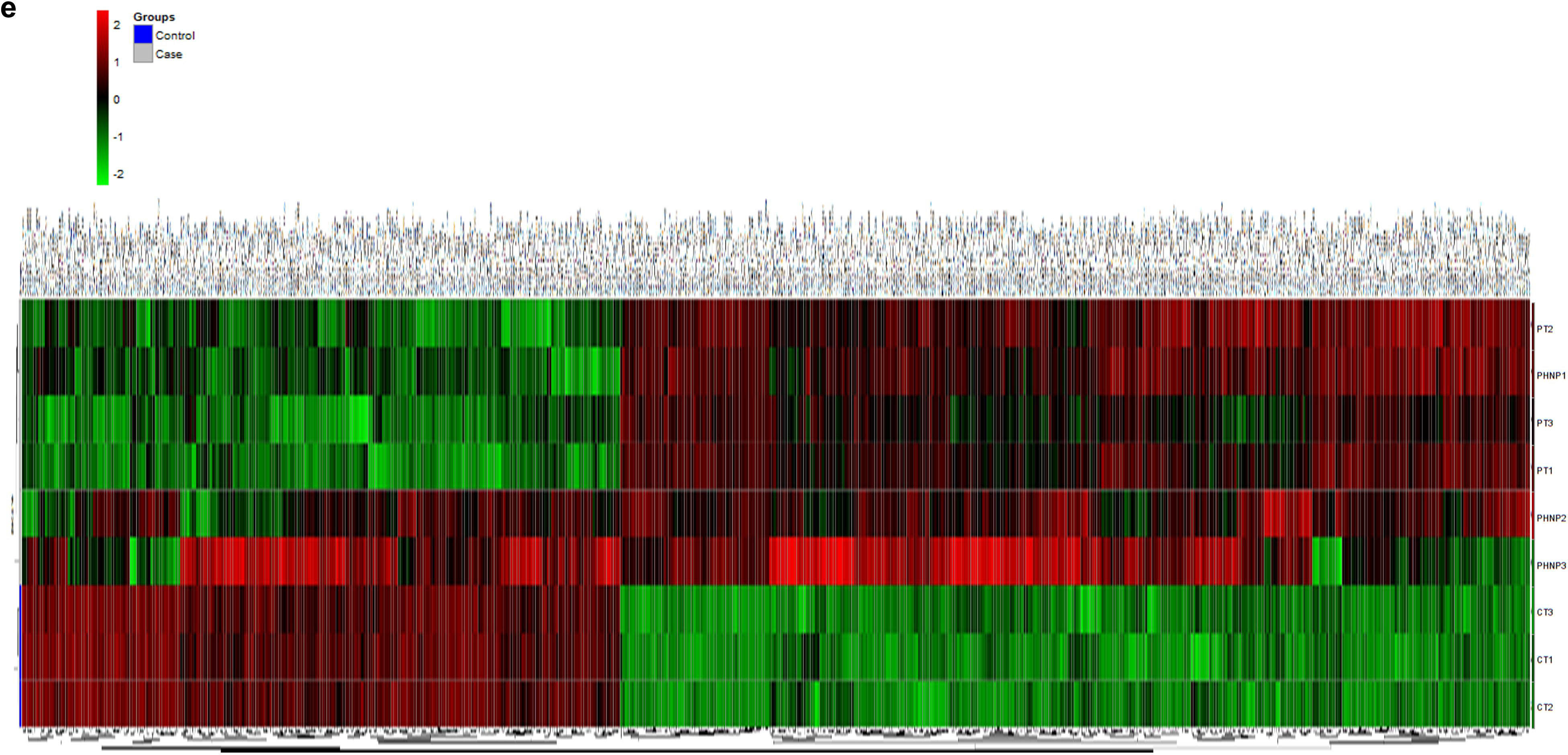
Differential expression of genes between different treated groups of *C. elegans*. **(a)** Volcano plot (p-value v/s log Fc) for PT group versus CT group. **(b)** scatter plot displaying the statistically significant differentially altered genes between PT treatment and control group (PT vs. CT). **(c)** Volcano plot (p-value v/s log Fc) for PHNP group versus PT group **(d)** scatter plot displaying the statistically significant differentially altered genes between peptide HNP pre-treated *C. elegans* followed by PT treatment and PT-treated *C. elegans*. CT: untreated worms, PT: PT treated worms, PHNP: custom peptide HNP pre-treatment followed by PT treatment, HNP: custom peptide HNP treated worms. **(e)** Heat map showing the differential expression of the upregulated and downregulated genes among different groups of *C. elegans*. PT: PT treated, PHNP: peptide HNP pre-treatment followed by PT treatment, CT: untreated (control) worms.

### 3.8. Quantitative mass spectrometry analysis demonstrated differential expression of cellular proteins between PT-treated and HNP peptide pre-treated followed by PT-treated *C. elegans*

A total number of 1026 non-redundant proteins were identified by the LC-MS/MS analysis from the different treatment groups of *C. elegans* (N2 strain). The identified proteins were classified into 20 discrete categories based on their cellular location and biological activity (Fig 9a, Supplementary Table S2), of which 13 major cellular proteins were found with high relative abundance.

**Fig 9.**
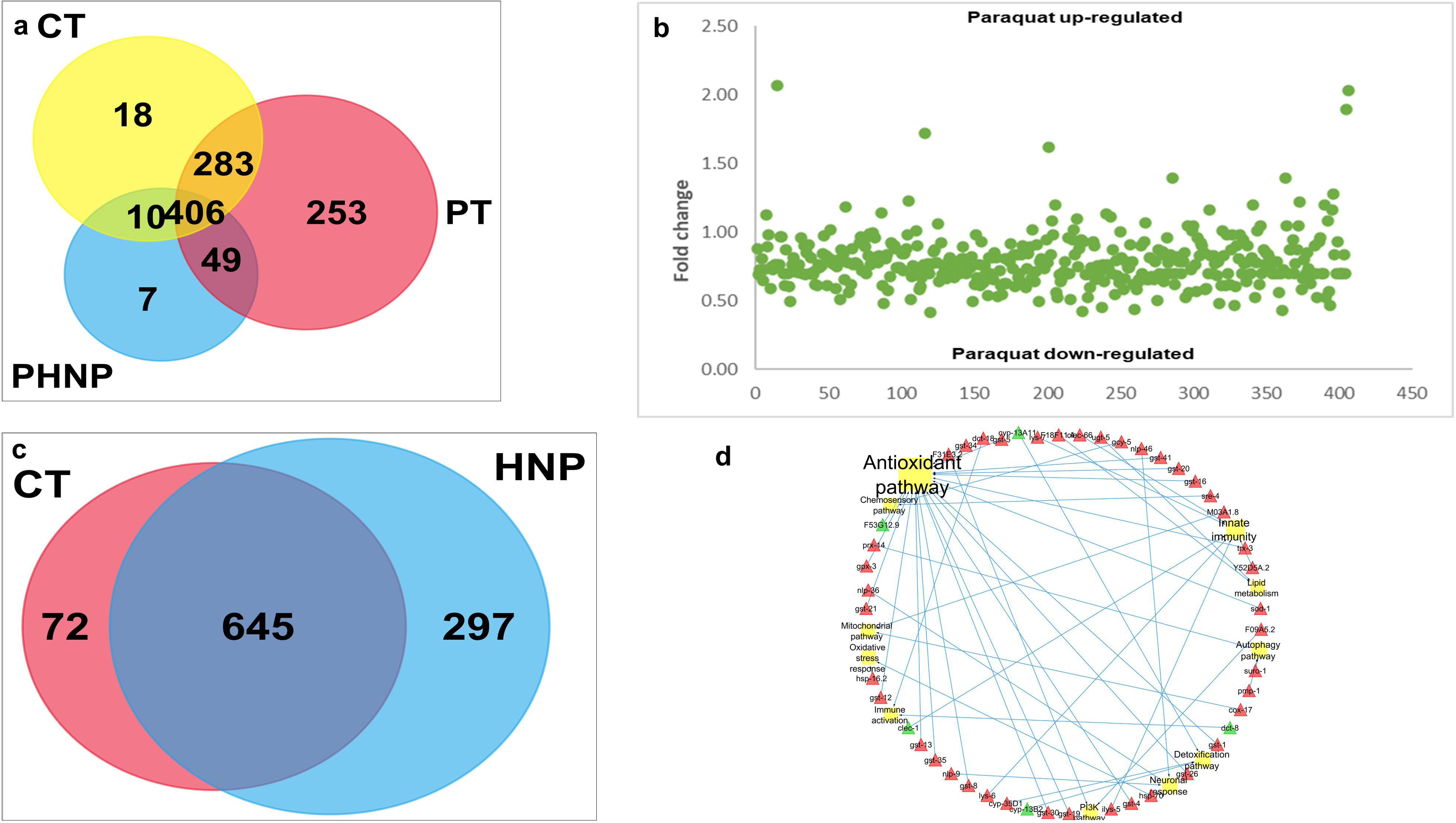
Proteomics analysis to show the expression of common and intracellular proteins among the treated groups of *C. elegans.* **(a)** Venn diagram showing common intracellular proteins among untreated (control) (CT), PT (PT) treated, and HNP pre-treated followed by PT-treated (PHNP) groups of *C. elegans* determined by LC/MS-MS analysis. **(b)** Scatter plot showing significantly upregulated (fold change>1.25) and downregulated (fold change <0.80) proteins in PT-treated *C. elegans*. FC: fold-change in expression determined by LC/MS-MS analysis. **(c)** Venn diagram showing common intracellular proteins among untreated (control) (CT) and only HNP-treated *C. elegans* determined by LC/MS-MS analysis. **(d)** Molecular network of custom peptide HNP-mediated neuroprotection. The interaction network of peptide HNP-regulated genes/proteins and interlinking pathways as determined by both transcriptomic and proteomic analyses.

In the PT group, 991 proteins were identified, among which differential expression was observed in 689 proteins compared to the control, and the expression of 406 proteins was restored in the PHNP group (Fig 9a). Between the PHNP and PT groups, 49 proteins were uniquely expressed (Fig 9a).

The scatter plot shows the differential expression of proteins in the PT group of *C*. *elegans* compared to the CT group of worms (Fig 9b). Further, the intracellular proteins of the PHNP group were compared with the PT and CT groups to identify the differentially expressed proteins involved in apoptosis, electron transport chain, stress response, antioxidant, and ubiquitin-proteasome pathways, which were restored in the PHNP group of worms (Supplementary Table S2). The intracellular protein of HNP-treated *C. elegans* was compared with the CT group, of which 297 proteins were uniquely expressed (Fig 9c). Proteomics analysis provided evidence of uniquely defined protein-regulated pathways in neuronal growth and development (Table 1).

**Table 1.** List of the uniquely expressed metabolic pathways in *C. elegans* (N2) treated with HNP compared to untreated (control) *C. elegans*. Quantitative proteomic analyses determined these pathways.

| Mapped ID/ Accession number | Pathways name | Proteins |
| --- | --- | --- |
| CAEEL\|WormBase=WBGene00006460\|UniProtKB=Q19775 | Activin beta signaling pathway | ppm-1.A |
| CAEEL\|WormBase=WBGene00004307\|UniProtKB=Q18246 | Heterotrimeric G-protein signaling pathway-Gi alpha and Gs alpha-mediated pathways | rap-1 |
| CAEEL\|WormBase=WBGene00004357\|UniProtKB=Q22038, CAEEL\|WormBase=WBGene00000390\|UniProtKB=Q05062 | Axon guidance mediated by Slit/Robo | rho-1 |
| CAEEL\|WormBase=WBGene00001748\|UniProtKB=P48727, CAEEL\|WormBase=WBGene00001747\|UniProtKB=Q27497 | Nicotine pharmacodynamics pathway | gsp-2 |
| CAEEL\|WormBase=WBGene00003951\|UniProtKB=Q9XUV0, CAEEL\|WormBase=WBGene00003949\|UniProtKB=Q23237 | Proteasome pathway | pbs-5 |
| CAEEL\|WormBase=WBGene00021826\|UniProtKB=G5EES9 | Oxidative stress response | txl-1 |
| CAEEL\|WormBase=WBGene00006460\|UniProtKB=Q19775 | ALP23B_signaling_pathway | ppm-1.A |
| CAEEL\|WormBase=WBGene00019599\|UniProtKB=O01590 | 5-Hydroxytryptamine degradation | acds-10 |
| CAEEL\|WormBase=WBGene00004460\|UniProtKB=Q04908 | Cell cycle | rpn-3 |
| CAEEL\|WormBase=WBGene00015337\|UniProtKB=P34277 | Vitamin C metabolism | gsto-2 |
| CAEEL\|WormBase=WBGene00020146\|UniProtKB=Q22067 | Neurogenesis | maph-1.1, |
| CAEEL\|WormBase=WBGene00001262\|UniProtKB=Q09590 | Vitamin D metabolism pathway | emb-8 |
| CAEEL\|WormBase=WBGene00001914\|UniProtKB=K7ZUH9, CAEEL\|WormBase=WBGene00003955\|UniProtKB=O02115 | DNA replication | pcn-1 |
| CAEEL\|WormBase=WBGene00006460\|UniProtKB=Q19775 | MYO_signaling_pathway | ppm-1.A |
| CAEEL\|WormBase=WBGene00008809\|UniProtKB=O17806, CAEEL\|WormBase=WBGene00010606\|UniProtKB=Q9XUT8 | Detoxification pathways | cyp-13B2, cyp-13A11 |
| CAEEL\|WormBase=WBGene00000547\|UniProtKB=Q9N4B1, CAEEL\|WormBase=WBGene00000390\|UniProtKB=Q05062, CAEEL\|WormBase=WBGene00000161\|UniProtKB=Q22601, CAEEL\|WormBase=WBGene00003171\|UniProtKB=P12456, CAEEL\|WormBase=WBGene00006537\|UniProtKB=P52275, CAEEL\|WormBase=WBGene00000183\|UniProtKB=G5EFK4, CAEEL\|WormBase=WBGene00001683\|UniProtKB=P04970 | Huntington disease | clp-7, cdc-42, mec-7, arf-3, gpd-1 |
| CAEEL\|WormBase=WBGene00001683\|UniProtKB=P04970 | Glycolysis | gpd-1 |
| CAEEL\|WormBase=WBGene00020557\|UniProtKB=P91455 | Adenine and hypoxanthine salvage pathway | T19B4.3 |
| CAEEL\|WormBase=WBGene00007175\|UniProtKB=Q17499 | Nicotinic acetylcholine receptor signalling pathway | B0395.3 |
| CAEEL\|WormBase=WBGene00021286\|UniProtKB=Q966C7 | Pentose phosphate pathway | tald-1 |
| CAEEL\|WormBase=WBGene00004357\|UniProtKB=Q22038,<br>CAEEL\|WormBase=WBGene00000390\|UniProtKB=Q05062,<br>CAEEL\|WormBase=WBGene00003171\|UniProtKB=P12456,<br>CAEEL\|WormBase=WBGene00006537\|UniProtKB=P52275,<br>CAEEL\|WormBase=WBGene00006794\|UniProtKB=Q07750 | Cytoskeletal regulation by Rho GTPase | rho-1,<br>cdc-42,<br>tbb-2,<br>unc-60 |
| CAEEL\|WormBase=WBGene00015064\|UniProtKB=Q09438 | Purine metabolism | B0228.7 |
| CAEEL\|WormBase=WBGene00012149\|UniProtKB=G5EDW8 | Allantoin degradation | CELE_VF13D12L.3 |
| CAEEL\|WormBase=WBGene00008654\|UniProtKB=Q19311,<br>CAEEL\|WormBase=WBGene00011064\|UniProtKB=Q21774 | De novo purine biosynthesis | pfas-1,<br>adsl-1 |
| CAEEL\|WormBase=WBGene00011745\|UniProtKB=Q94048,<br>CAEEL\|WormBase=WBGene00001532\|UniProtKB=Q23682,<br>CAEEL\|WormBase=WBGene00009306\|UniProtKB=P91859 | Chemosensory response | sre-4, gcy-5,<br>got-1.2 |
| CAEEL\|WormBase=WBGene00017772\|UniProtKB=Q22966, | Innate immune response pathways | clec-1 |
| CAEEL\|WormBase=WBGene00000390\|UniProtKB=Q05062 | TGF-beta signalling pathway | cdc-42 |
| CAEEL\|WormBase=WBGene00003951\|UniProtKB=Q9XUV0,<br>CAEEL\|WormBase=WBGene00004505\|UniProtKB=O76371,<br>CAEEL\|WormBase=WBGene00004501\|UniProtKB=Q18787,<br>CAEEL\|WormBase=WBGene00004460\|UniProtKB=Q04908 | Ubiquitin proteasome pathway | pbs-5,<br>rpt-5, |
| CAEEL\|WormBase=WBGene00003747\|UniProtKB=H2KZL2<br>CAEEL\|WormBase=WBGene00012145\|UniProtKB=Q7YTJ1<br>CAEEL\|WormBase=WBGene00007185\|UniProtKB=Q03561 | Neuronal process | nlp-36<br>nlp-9<br>nlp-46 |
| CAEEL\|WormBase=WBGene00006460\|UniProtKB=Q19775 | GBB_signaling_pathway | ppm-1.A |
| CAEEL\|WormBase=WBGene00012149\|UniProtKB=G5EDW8 | TCA cycle | CELE_VF13D12L.3 |
| CAEEL\|WormBase=WBGene00007175\|UniProtKB=Q17499 | Muscarinic acetylcholine receptor 1 and 3 signalling<br>Pathway | B0395.3 |
| CAEEL\|WormBase=WBGene00012149\|UniProtKB=G5EDW8,<br>CAEEL\|WormBase=WBGene00014001\|UniProtKB=Q23539,<br>CAEEL\|WormBase=WBGene00020139\|UniProtKB=P91408 | Pyruvate metabolism | pyk-2,<br>eppl-1 |

### 3.9. Transcriptomic and functional proteomics analyses in unison have elucidated the reversal of PT-induced upregulated and downregulated metabolic pathway genes by pre-treatment of *C*. *elegans* with neuroprotective peptide

The transcriptomic and quantitative proteomics data explicitly demonstrated the regulation of commonly upregulated and downregulated genes and their intracellular proteins in the PT and HNP groups compared to the CT group of *C. elegans* (N2 strain). The upregulated and downregulated proteins are involved in apoptosis, superoxide removal, lipid and protein metabolism, cellular differentiation, energy metabolism, mitochondrial function, axonal transport, autophagy, and neuronal development pathways in *C. elegans*. The analyses showed that the 15 upregulated (Table 2a) and 25 downregulated (Table 2b) signaling pathways in PT-treated *C. elegans* (N2 strain) were restored to an average level by the HNP peptide pre-treatment before the addition of PT (Tables 2a,b).

**Table 2a.** Transcriptomic and proteomic analyses to show the paraquat-induced upregulated metabolic pathways in wild type N2 strain of *C. elegans*. These pathways were restored to control (normal) level by pre-treatment of worms with neuroprotective custom peptide HNP.

| Accession Number | Mapped ID | Metabolic pathway name |
| --- | --- | --- |
| P02748 | CAEEL\|WormBase=WBGene00001149\|UniProtKB=P54688,<br>CAEEL\|WormBase=WBGene00020831\|UniProtKB=O61856 | Isoleucine biosynthesis |
| P00027 | CAEEL\|WormBase=WBGene00001648\|UniProtKB=P51875 | Heterotrimeric G-protein signalling pathway-Gq alpha<br>and Go alpha-mediated pathway |
| P06959 | CAEEL\|WormBase=WBGene00002183\|UniProtKB=Q22100 | CCKR signalling map |
| P00053 | CAEEL\|WormBase=WBGene00000552\|UniProtKB=O16305 | T cell activation |
| P00055 | CAEEL\|WormBase=WBGene00002190\|UniProtKB=P30625 | Transcription regulation by bZIP transcription factor |
| P00059 | CAEEL\|WormBase=WBGene00002363\|UniProtKB=G5EGK8 | p53 pathway |
| P02723 | CAEEL\|WormBase=WBGene00010083\|UniProtKB=O17892 | Adenine and hypoxanthine salvage pathway |
| P05912 | CAEEL\|WormBase=WBGene00002190\|UniProtKB=P30625 | Dopamine receptor-mediated signalling pathway |
| P00006 | CAEEL\|WormBase=WBGene00017121\|UniProtKB=P19974 | Apoptosis signalling pathway |
| P00021 | CAEEL\|WormBase=WBGene00002363\|UniProtKB=G5EGK8 | FGF signalling pathway |
| P00034 | CAEEL\|WormBase=WBGene00000942\|UniProtKB=P19826,<br>CAEEL\|WormBase=WBGene00016913\|UniProtKB=Q18823 | Integrin signalling pathway |
| P02724 | CAEEL\|WormBase=WBGene00001149\|UniProtKB=P54688 | Alanine biosynthesis |
| P00030 | CAEEL\|WormBase=WBGene00006922\|UniProtKB=Q19213 | Hypoxia response via HIF activation |
| P04393 | CAEEL\|WormBase=WBGene00016236\|UniProtKB=Q18313,<br>CAEEL\|WormBase=WBGene00021022\|UniProtKB=Q9UA62 | Ras Pathway |
| P02745 | CAEEL\|WormBase=WBGene00014095\|UniProtKB=Q23621 | Glutamine glutamate conversion |

**Table 2b.** Transcriptomic and proteomics analyses to show the paraquat-induced downregulated metabolic pathways in N2 strain *C. elegans*. These pathways were restored to normal (control) levels by pre-treatment of worms with neuroprotective custom peptide HNP.

| Accession Number | Mapped ID | Metabolic pathway name |
| --- | --- | --- |
| P00012 | CAEEL\|WormBase=WBGene00008809\|UniProtKB=O17806,<br>CAEEL\|WormBase=WBGene00010606\|UniProtKB=Q9XUT8 | Detoxification pathways |
| P02746 | CAEEL\|WormBase=WBGene00010645\|UniProtKB=G5EBW5,<br>CAEEL\|WormBase=WBGene00001337\|UniProtKB=Q23315 | Heme biosynthesis |
| P00018 | CAEEL\|WormBase=WBGene00001515\|UniProtKB=Q22720,<br>CAEEL\|WormBase=WBGene00004287\|UniProtKB=Q94124,<br>CAEEL\|WormBase=WBGene00002335\|UniProtKB=P22981 | EGF receptor signalling pathway |
| P00019 | CAEEL\|WormBase=WBGene00013785\|UniProtKB=Q9U2T1,<br>CAEEL\|WormBase=WBGene00013786\|UniProtKB=Q9U2T0 | Endothelin signalling pathway |
| P02771 | CAEEL\|WormBase=WBGene00016103\|UniProtKB=Q18164 | Pyrimidine metabolism |
| P00004 | CAEEL\|WormBase=WBGene00011745\|UniProtKB=Q94048,<br>CAEEL\|WormBase=WBGene00001532\|UniProtKB=Q23682,<br>CAEEL\|WormBase=WBGene00009306\|UniProtKB=P91859 | Chemosensory response |
| P00048 | CAEEL\|WormBase=WBGene00012816\|UniProtKB=Q9XWX3,<br>CAEEL\|WormBase=WBGene00017183\|UniProtKB=O16575,<br>CAEEL\|WormBase=WBGene00002335\|UniProtKB=P22981 | PI3 kinase pathway |
| P00060 | CAEEL\|WormBase=WBGene00004466\|UniProtKB=O61742,<br>CAEEL\|WormBase=WBGene00004503\|UniProtKB=P46502,<br>CAEEL\|WormBase=WBGene00004465\|UniProtKB=Q22253 | Ubiquitin proteasome pathway |
| P00052 | CAEEL\|WormBase=WBGene00002335\|UniProtKB=P22981 | TGF-beta signalling pathway |
| P00049 | CAEEL\|WormBase=WBGene00020146\|UniProtKB=Q22067 | Neurogenesis |
| P00013 | CAEEL\|WormBase=WBGene00004466\|UniProtKB=O61742,<br>CAEEL\|WormBase=WBGene00004465\|UniProtKB=Q22253 | Cell cycle |
| P00031 | CAEEL\|WormBase=WBGene00003747\|UniProtKB=H2KZL2<br>CAEEL\|WormBase=WBGene00012145\|UniProtKB=Q7YTJ1<br>CAEEL\|WormBase=WBGene00007185\|UniProtKB=Q03561 | Neuronal process |
| P00016 | CAEEL\|WormBase=WBGene00003171\|UniProtKB=P12456,<br>CAEEL\|WormBase=WBGene00004287\|UniProtKB=Q94124 | Cytoskeletal regulation by Rho GTPase |
| P00021 | CAEEL\|WormBase=WBGene00001515\|UniProtKB=Q22720,<br>CAEEL\|WormBase=WBGene00004287\|UniProtKB=Q94124,<br>CAEEL\|WormBase=WBGene00006352\|UniProtKB=G5EDR3,<br>CAEEL\|WormBase=WBGene00002335\|UniProtKB=P22981 | FGF signalling pathway |
| P00044 | CAEEL\|WormBase=WBGene00012437\|UniProtKB=Q9XWR0,<br>CAEEL\|WormBase=WBGene00002348\|UniProtKB=P02567,<br>CAEEL\|WormBase=WBGene00003514\|UniProtKB=P12845 | Nicotinic acetylcholine receptor signalling pathway |
| P02721 | CAEEL\|WormBase=WBGene00010419\|UniProtKB=Q9XXK1,<br>CAEEL\|WormBase=WBGene00000229\|UniProtKB=P46561,<br>CAEEL\|WormBase=WBGene00022089\|UniProtKB=Q95XJ0 | ATP synthesis |
| P00009 | CAEEL\|WormBase=WBGene00004287\|UniProtKB=Q94124 | Axon guidance mediated by netrin |
| P00008 | CAEEL\|WormBase=WBGene00004287\|UniProtKB=Q94124,<br>CAEEL\|WormBase=WBGene00004357\|UniProtKB=Q22038,<br>CAEEL\|WormBase=WBGene00000390\|UniProtKB=Q05062 | Axon guidance mediated by Slit/Robo |
| P00045 | CAEEL\|WormBase=WBGene00000075\|UniProtKB=Q94316 | Notch signalling pathway |
| P00007 | CAEEL\|WormBase=WBGene00016104\|UniProtKB=Q18163 | Axon guidance mediated by semaphorins |
| P00051 | CAEEL\|WormBase=WBGene00018491\|UniProtKB=Q9UAV5,<br>CAEEL\|WormBase=WBGene00001503\|UniProtKB=O17214,<br>CAEEL\|WormBase=WBGene00007350\|UniProtKB=P53596,<br>CAEEL\|WormBase=WBGene00020679\|UniProtKB=O61199,<br>CAEEL\|WormBase=WBGene00000041\|UniProtKB=P34455,<br>CAEEL\|WormBase=WBGene00000833\|UniProtKB=P34575 | TCA cycle |
| P00017 | CAEEL\|WormBase=WBGene00008645\|UniProtKB=P90829,<br>CAEEL\|WormBase=WBGene00004340\|UniProtKB=P53016 | DNA replication |
| P00046 | CAEEL\|WormBase=WBGene00015879\|UniProtKB=O45155 | Oxidative stress response |
| P00009 | CAEEL\|WormBase=WBGene00000390\|UniProtKB=Q05062 | Axon guidance mediated by netrin |
| P00024 | CAEEL\|WormBase=WBGene00011884\|UniProtKB=Q27527,<br>CAEEL\|WormBase=WBGene00011474\|UniProtKB=P54216,<br>CAEEL\|WormBase=WBGene00006601\|UniProtKB=Q10657,<br>CAEEL\|WormBase=WBGene00020185\|UniProtKB=P91427 | Glycolysis |

The comparative data obtained from transcriptomic, proteomics, and qRT-PCR studies suggest that PT treatment induced upregulation of apoptotic pathways, heterotrimeric G-protein signaling pathway-Gq alpha and Go alpha mediated pathway, ras pathway, p53 pathway, T-cell activation pathway, dopamine (DA) receptor-mediated signaling pathway, and hypoxia response via HIF activation in *C. elegans* (Table 2a). In contrast, the downregulation of essential pathways such as antioxidative, detoxification, chemosensory response, EGF receptor signaling, PI3K, ubiquitin-proteasome, neurogenesis, TGF-beta signaling, Axon guidance, FGF signaling, nicotinic acetylcholine receptor signaling, ATP synthesis, TCA cycle, DNA replication, oxidative stress response, glycolysis, and neuronal development was noted post-PT-treatment in *C*. *elegans* (Table 2b). Likewise, the collective analysis of transcriptomic and quantitative proteomics data showed that the HNP group, compared to the CT group, induced the expression of 33 unique signaling pathways (Table 1). Moreover, overexpression of the metabolic pathway genes (Table 2b) in the PHNP group explicitly validated the enhanced biosynthesis of biomolecules required for neurogenesis and neuroprotection mechanism against PT-induced toxicity in *C. elegans*.

### 3.10. The molecular network analysis demonstrates the neuroprotective functions of HNP against PT-induced toxicity and neuronal damage

Transcriptomic and proteomic analyses revealed different metabolic pathways involved in HNP-induced protection against PT-induced neuronal damage in N2 strain *C. elegans*. Based on these pieces of information, the molecular interactions, a network of gene expression profiles was constructed (Fig 9d). The mapping showed that the interconnected pathways of PT-induced knock-over genes leading to neuronal dysfunction were restored with the HNP pre-treatment in *C. elegans* (Fig 9d). In unison, transcriptomic and proteomic studies have identified 15 common upregulated pathways between the PHNP and PT groups of *C. elegans.* The molecular network displayed the snapshot of the genes regulating proteins and their biological functions, modulating the neurogenesis and neuroprotective function of HNP against PT-induced toxicity and neuronal dysfunction (Fig 9d, Table 1).

An overall neuroprotective mechanism against PT-induced neurotoxicity is proposed in Fig 10

**Fig 10.**
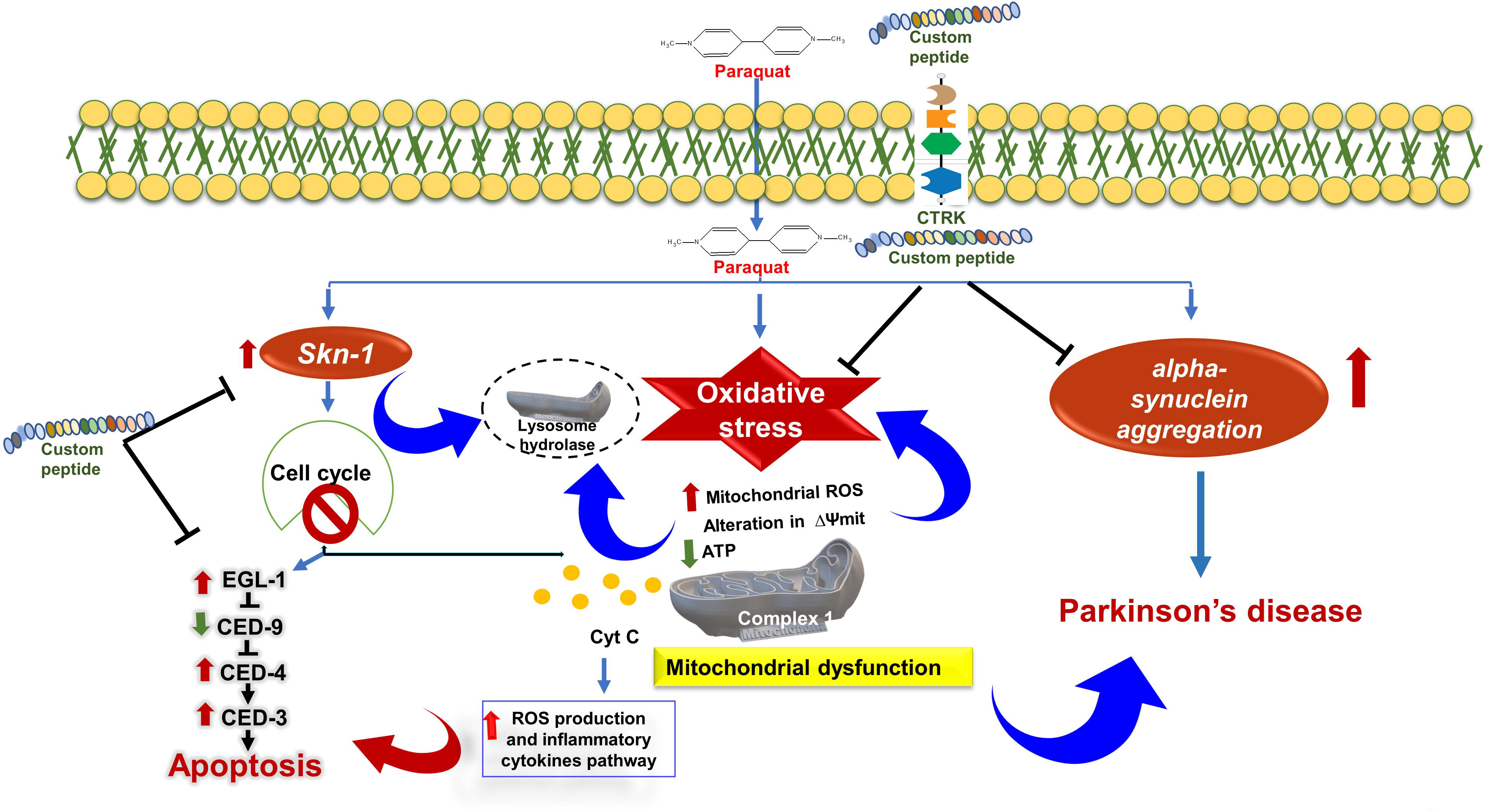
The proposed neuroprotection mechanism pathways of custom peptide mediated protection against PT-induced neurotoxicity in *C. elegans* (N2 strain). DAergic neuronal degeneration and α-synuclein aggregation are two major hallmarks of PD. The PT-induced overexpression of SKN-1 (p38/MAPK) induced response does not protect DAergic neurons from degeneration, affect cell cycle and trigger the activation of apoptotic pathways. Upregulation of antioxidant enzymes (sods and catalases), aggregation of ROS, and mitochondrial release of cytochrome c upon exposure to neurotoxins such as PT is one of the crucial factors in DAergic neuron degeneration, α-synuclein aggregation, and neuronal death. Custom peptide-pretreatment downregulates SKN-1 pathways, increase stress resistance, ameliorates mitochondrial stress and inhibit activation of apoptotic pathways. Custom peptides-mediated upregulation of the *cat-2* gene regulates tyrosine hydroxylase (TH) enzyme expression and restores the PT-induced DA deficiency.

### 3.11. Custom peptides were non-toxic to mice and safe to administer

Custom peptides (TNP and HNP at the ratio 1:1, 10.0 mg/kg) were non-toxic to mice and did not show behavioral changes or adverse effects in treated mice. The serum profiles viz. SGOT, ALKP, SGPT, BUN, glucose content, creatinine, cholesterol, bilirubin and albumin of the custom peptide-treated mice (24 h post-treatment) did not show any significant (p ≧ 0.05) deviation compared to the control (1X PBS treated) group of mice (Supplementary Table S3). A minor increase in glucose level was measured in the blood of the treated mice compared to control mice, but it was within the normal range of blood glucose in mice (Supplementary Table S3). Moreover, microscopic examination of the brain, heart, kidney, liver, lung, ovary, and testis of the custom peptide (HNP)-treated mice showed no morphological alterations (Supplementary Fig S9). However, HNP treatment significantly (p ≤ 0.05) reduced the production of proinflammatory cytokines viz. TNF-α, IL-6, IL-1β, compared to the control group of mice (Supplementary Fig S10).

## 4. Discussion

In recent years, the characterization of novel neurotrophic peptides has gained more focus in clinical trials against different NDs [23, 28, 48–50]. superior neuroprotective effect, synaptic and neural plasticity, neurogenesis, superior pharmacokinetics than the parent neurotrophin, and easier penetration are only some benefits that neurotrophic peptides offer over other neurotrophic medications [8, 51].

Our previous study has demonstrated the exclusive binding of TNP and HNP to the TrkA receptor of PC-12 cells to induce neuritogenesis [52]; however, *C. elegans* possesses a simple nervous system and lacks Trk receptors [53]. Therefore, we were puzzled to understand the region of binding and mechanism of neuroprotective activity of custom peptides in *C. elegans*. To solve this issue, we investigated the presence of TrkA receptor homolog in the *C. elegans*. The presence of TrkA receptor homolog CAM-1 receptor in *C. elegans* was discovered by BLAST search.

Further *in silico* studies validated the binding of custom peptides to the CAM-1 receptor. The CAM-1 in *C. elegans* is a less functionally explored protein receptor. However, some reports suggest that the CAM-1 is involved in essential functions, viz., the regulation of asymmetric division of V cells (seam cells), CA/CP neuroblast, and axon outgrowth [54, 55] and the positioning of the nerve ring [56]. Further, the CAM-1 receptor is also responsible for the negative regulation of developmental neurite pruning of AIM neurons [57]. Moreover, Entries in the Reactome pathway browser database [45] demonstrated that the CAM-1 receptor is an essential component in the MAPK signalling pathway by playing multiple roles, such as the regulation and development of axon [54, 55] and the positioning of the nerve ring [56].

Therefore, the CAM-1 receptor as a confirmed binding site for the custom peptides in this study was further validated by insignificant binding of FITC-custom peptides in the CAM-1 mutant strain of *C. elegans* at its nerve ring region. However, FITC-conjugated custom peptides showed binding to *C. elegans* at its nerve ring adjacent to the pharynx, which covers complex circuitry leading to most aspects of behaviors, such as chemo-sensation, locomotion, learning, memory, etc. [58]. The nerve ring of *C. elegans* adjacent to the pharynx functions similarly to a ’’brain,’ and most sensory neurons have endings organized around the mouth [58].

PD is associated with multiple motor and non-motor symptoms developed due to elevated oxidative stress leading to mitochondrial membrane damage, which triggers DAergic neuron degeneration [59]. A plethora of reports support that oxidative stress is one of the essential factors associated with the progression of NDs such as PD, and evidence has been presented to show that antioxidant activity may prevent ND [60, 61]. We have demonstrated the *in vitro* antioxidant properties of the TNP and HNP by their free radical scavenging activity [20]. Therefore, TNP and HNP *in vivo* conditions also prolong lifespans by decreasing oxidative stress in worms [62, 63]. Antioxidants such as vitamin C and quercetin demonstrated neuroprotection properties in several *in vitro* and *in vivo* models of NDs viz. AD, PD, and Huntington’s Disease (HD) [64–66]. Therefore, we considered quercetin and vitamin C a positive control for this study.

Since ASH sensory neurons of *C. elegans* play a pivotal role in the sensory response to aversive stimuli, PT treatment induces defective chemotaxis behavior in worms due to the loss of ASH neuronal function [47]. Therefore, the chemotaxis assay was performed in *C. elegans* to demonstrate the possibility of restoration of PT-induced chemotaxis dysfunction by pre-treatment with custom peptides. The pre-treatment of *C. elegans* with custom peptides before PT treatment has shown a superior protective effect by restoring the PT-induced chemotaxis defect and survivability, which may be correlated to the fact that these peptides bind to the same region (nerve ring adjacent to the pharynx) where PT binds to induce neurodegeneration in *C. elegans*. Therefore, post-administration of peptides after PT treatment, minimal protection was observed against PT-induced neurodegeneration when they already bind to neurons. Further, peptides’ neuroprotective and neurotrophic effects were validated from the transcriptomic and proteomic analysis data. Both unambiguously showed that the proteins responsible for chemosensory behavior, viz. sre-4, gcy-5, and got-1.2 were downregulated by PT, resulting in the loss of memory of *C*. *elegans*. In sharp contrast, pre-treatment with custom peptides upregulated these genes to increase the cellular proteins to an average level (untreated control worms).

The increased ROS content observed in the neurons of PD patients and PD-like model organisms triggers cellular damage and apoptosis [63, 67, 68]. Therefore, antioxidant compounds, for example, quercetin, curcumin, naringin, metformin, bacosides, hydralazine, and vitamin C, have found significant therapeutic potential to treat PT-induced toxicity [24, 69–71]. For averting non-autonomous neurodegeneration and cellular damage, mitochondrial failure sets off a cascade of stress response pathways, including the innate immune pathways p38/MAPK [72]. HNP and TNP (50 µg/mL, [41.6 µM (TNP) and 35.7 µM (HNP)]) pre-treatment for 2 h demonstrated appreciable potency to reduce the PT-mediated increase in cellular ROS level and mitochondrial membrane depolarization, subsequently enhancing survival rates in PT-treated *C. elegans*. Vitamin C pre-treatment for 24 h significantly attenuated the PT-induced ROS production but did not restore mitochondrial membrane depolarization at a concentration (100 µg/mL, 568 µM) two times higher than the concentration of custom peptides. Furthermore, vitamin C could not protect against PT-induced neurotoxicity, suggesting that the peptides’ antioxidant properties may not be the primary means of preventing PT-induced neuronal damage. This study supports the therapeutic efficacy of the peptides under investigation.

In response to oxidative stress, the *skn-1* protein translocate and accumulates at the intestinal nuclei and induces transcription of genes such as SODs (*sod-1* and *sod-3*), catalases (*cat-1*, *cat-2* and *cat3*), thioredoxin-1 (*trx-1*), GSTs (*gst-4*, *gst-6* and *gst-10*), and heat-shock proteins HSPs (*hsp-16.1* and *hsp-16.2*) involved in phase 2 detoxification [38]. Accordingly, the MAPK genes *sek-1*, *pmk-1*, and *skn-1* were upregulated in *C. elegans* post-PT exposure. However, the qRT-PCR results have shown the overexpression of downstream genes of activated p38/MAPK pathways such as ’*skn-‘1’* that regulates various detoxification processes (*sod*, *cat*, *trx*, *gst* series) and heat-shock proteins (HSPs) genes essential to abolish enhanced ROS production leading to cellular damage [72, 73]. The expression level of SODs and other associated genes is low under normal conditions [37, 73]. However, acute exposure to PT induces higher expression of such genes in mitochondria to withstand oxidative stress. The PT-mediated over-expression of HSPs prevents the protein from misfolding due to changes in the cellular redox state [74]. The qRT-PCR result highlighted the significant downregulation of stress-related genes in custom peptide-treated wild-type N2 worms compared to the PT treatment, probably due to the enhanced transcription initiation or mRNA stability. Downregulation of these genes upon custom peptide pre-treatment is correlated with reduced oxidative stress and increased stress resistance in *C. elegans*, which concludes the antioxidant and neuroprotective potential of TNP and HNP.

DAergic neurons are vulnerable to PT-induced oxidative stress due to ROS generation, a principal neuronal death regulator [75]. DAergic neuronal degeneration and α-synuclein aggregation are two major hallmarks of PD [76]. The PT-induced overexpression of SKN-1 (p38/MAPK) induced response does not protect DAergic neurons from degeneration [72]. Upregulation of antioxidant enzymes (sods and catalases) and aggregation of ROS upon exposure to neurotoxins such as PT is one of the crucial factors in DAergic neuron degeneration and α-synuclein aggregation [77]. Custom peptides-mediated upregulation of the *cat-2* gene regulates tyrosine hydroxylase (TH) enzyme expression and restores the PT-induced DA deficiency [41] (Fig 10).

The neurotransmitter DA availability to the striatum (brain structure) is essential for motor response and memory functions [78, 79]. DA deficiency is another hallmark of PD, resulting in DAergic cell death and α-synuclein deposition [79, 80]. The neuroprotective activity of custom peptides against PT-induced chemotaxis behavior defects, DAergic neurodegeneration, α-synuclein aggregation, and reduced life span may be correlated to their antioxidant and antiapoptotic properties [75]. However, vitamin C was ineffective in restoring chemotaxis behavior, DAergic neurodegeneration, and α-synuclein aggregation induced by the acute exposure of PT in *C. elegans*, so reinforcing antioxidation may not be the only property to offer protection against PT-induced neurodegeneration [20].

Because the unique peptide HNP prevented PT-induced toxicity better than TNP, it was chosen for additional research. Compiling the transcriptome, proteomics, and qRT-PCR research results, custom peptides pre-treatment unambiguously showed that PT-induced elevated pro-apoptotic (cytochrome C, CED-4, and CED-3) gene expression was downregulated. On the other hand, it increased the expression of antiapoptotic genes (CED-9) in *C. elegans* treated with PT, increasing the worms’ survival rate.

The transcriptomic and quantitative proteomic analyses have identified the molecular network of signaling pathways through which the peptide HNP blocked the PT-induced neurotoxicity in *C. elegans*. The findings aided that HNP pre-treatment restored the PT-induced altered genes and protein functions such as anti-oxidant pathways, mitochondrial stress response pathways, energy metabolism pathways, lipid and protein metabolism pathways, and programmed cell death in *C. elegans*.

The custom peptide also restored altered pathways induced by the PT treatment in the *C. elegans* PD model (N2 strain). PT causes its toxic effect by upregulating most of the antioxidant genes (sod, trx, gst), mitochondrial stress resistance gene (txl-1, cox-4), detoxification genes (cyp-13B2, cyp-13A11), TCA cycle (CELE_VF13D12L), genes involved in chemosensory response (sre-4, gcy-5, got-1.2), neurogenesis (maph-1.1), neuronal development (nlp-36, nlp-9, nlp-46) and upregulated apoptotic genes (cyt C, egl-1), MAPK genes involved in innate immune response (skn-1), and some of the stress response genes (hsp10) in *C. elegans*, which were counteracted by custom peptide (HNP) pre-treatment.

In summary, compared to the CT group of C. elegans, the PT group of worms showed the most significant variance in differential gene expression. On the other hand, the pre-treatment group of bespoke peptides displayed a minimum variance compared to the CT group, indicating the restoration or normalization of aberrant dynamics of genes and proteins. This work emphasizes the therapeutic value of tailored peptides in reducing neurodegeneration.

Using the tested custom peptides as a drug prototype exclusively depends upon their non-toxic nature in preclinical studies. The acute toxicity studies in mice models showed that the peptide is devoid of toxicity in mice at a dose that is 100 times higher than its therapeutic dose determined in *C. elegans*. Further, it has no detrimental effect on the biochemical parameters of blood parameters and vital organs, suggesting the safety of the peptides for the development of drug prototypes. Furthermore, the reduced concentration of inflammatory mediators (TNF-α, IL-6, IL-1β) in the HNP-treated mice, compared to control (1X PBS-treated) mice, diminished the risk of inflammatory response post-treatment with peptides.

One of our future objectives is to evaluate the pharmacokinetics, pharmacodynamics, and neuroprotective effect of the customized peptides in rodent models. Additionally, investigations on transcriptomics and proteomics have offered a comprehensive understanding of the general pathways that tailored peptides up or down-regulate to guard against PT-induced neurodegeneration. A more thorough understanding of the mechanisms underlying custom peptides is necessary.

## 5. Conclusion

The development of effective and safe drugs against neurodegenerative disorders such as ’ ’PD is still a challenging task. Our previous study demonstrated that custom peptides had the potential to protect and restore PT-mediated counteract PT-induced neurotoxicity by thwarting excessive ROS production, oxidative stress, MMP, and premature apoptotic death in PC-12 cells. In this study, we have validated our findings in a higher in vivo *C. elegans* model. Moreover, the present study also demonstrated that custom peptides counteract the DAergic neuron degeneration in BZ555 worms, restored loss of chemotaxis behavior, upregulated ROS, restored altered MMP, and prevented apoptotic cell death in N2 worms. Custom peptides also diminished α-synuclein aggregation in NL5901 worms. Moreover, qRT-PCR, transcriptomics, and proteomics studies unveiled the broad mechanism of neuroprotective activity of custom peptides in the PT-induced PD model of *C. elegans* N2 worms. However, preclinical studies of custom peptides in mice models are warranted to provide insights for developing safe therapeutic molecules for against NDs. Future in-depth studies are required to evaluate the pharmacokinetics, pharmacodynamics, and neuroprotective effect of the customized peptides in rodent models.

## Acknowledgments

The authors thank C-CAMP, NCBS Bangalore, India, for LC-MS/MS analysis, Biokart India Pvt Ltd, Bangalore, India, for transcriptomics analysis, and the Sophisticated Analytical Instrumentation Centre (SAIC), IASST for instrument facility. DM received SRF from the SERB project, and AP received a postdoctoral fellowship from IASST. The authors also thank This work was funded by the Science and Engineering Research Board, New Delhi (EMR/2017/001829) to AKM.

## Author’s Contribution

AKM conceived the idea, designed the study, and corrected and edited the manuscript; DM performed the experiments, analyzed the data, and wrote the manuscript; AP analyzed the proteomics and transcriptomics data, corrected the manuscript; RM performed experiments with CAM-1 mutant strain; KS and PAF performed the in silico data analysis; AK helped in the qRT-PCR analysis; MRK analyzed the data and corrected the manuscript.

## Data availability

All experimental raw data and custom scripts supporting this study’s findings are available from the corresponding author upon request.

## Conflict of interest

The authors have no relevant financial or non-financial interests to disclose.

## Ethics approvals

Ethical permission was approved by the animal ethics committee of IASST (IASST/IAEC/2022/10).

## Abbreviations

The abbreviations used are

CGC: Caenorhabditis Genetics Center
HNP: heptadeca-neuropeptide
NGF: nerve growth factor
NGM: nematode growth medium
PT: paraquat
qRT-PCR: quantitative reverse transcription-polymerase chain reaction
TNP: trideca-neuropeptide
TrkA: tropomyosin receptor kinase A receptor.

## Notes

### Competing Interest Statement

The authors have declared no competing interest.

