## Supplementary Table and Figure for "Snake venom-inspired novel peptides protect *Caenorhabditis elegans* against paraquat-induced Parkinson’s pathology"

**Running title:** Neuroprotective peptides against neurodegeneration.

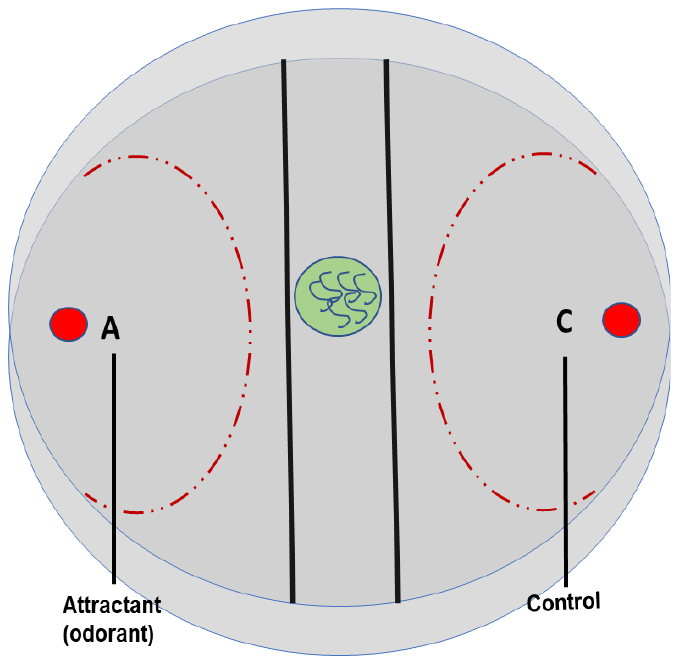

**Supplementary Fig S1.** The schematic diagram represents the experimental design of the chemotaxis assay to calculate the chemotaxis index (CI).

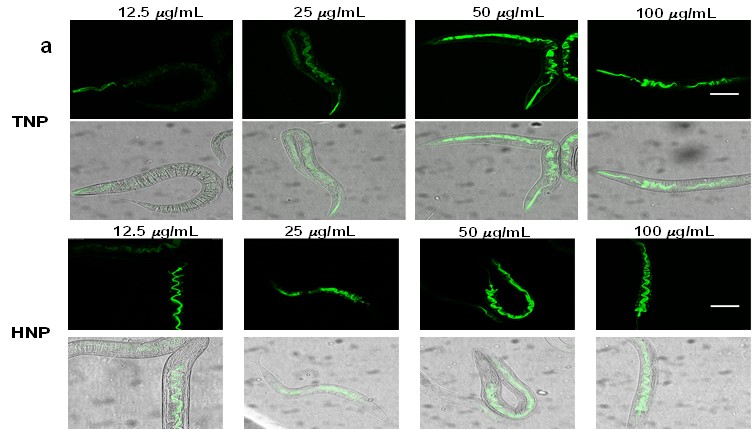

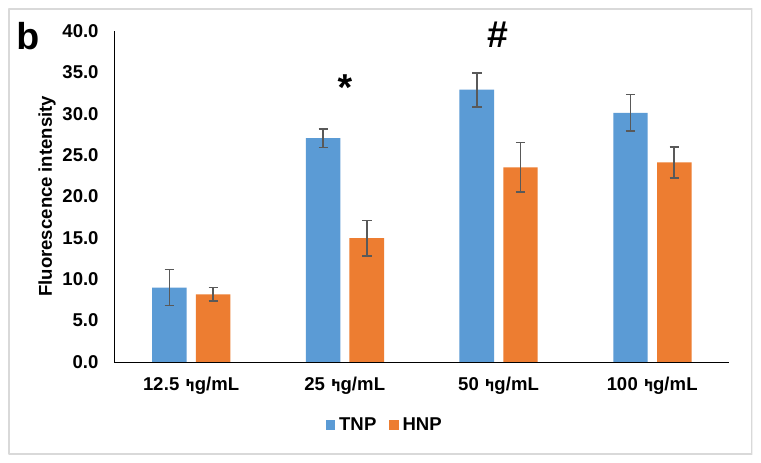

**Supplementary Fig S2.** Confocal microscopic (40 X) studies of the *in-vivo* binding of FITC-custom peptides to *C. elegans* for 2h. **(a)** Dose-dependent (12.5 µg/mL – 100 µg/mL) binding of custom peptides to the *C. elegans*. The scale bar indicates the length as 100 µm**. (b)** Bar graph representing fluorescence intensity between the treatment groups; ^*^p ≤ 0.05, a significant difference between 12.5 µg/mL and 25 µg/mL dose of FITC-conjugated peptides to *C. elegans*, ^#^p ≤ 0.05, a significant difference between 25 µg/mL and 50 µg/mL dose of FITC-conjugated peptides to *C. elegans*. Values are mean ± SD of triplicate determinations.

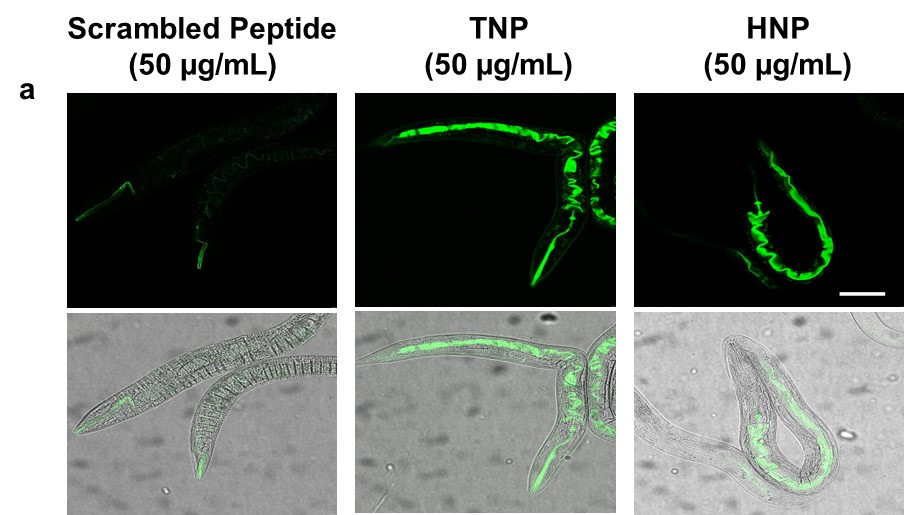

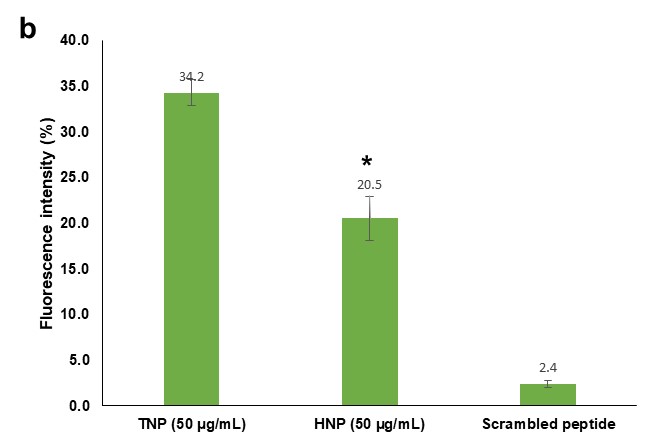

**Supplementary Fig S3.** Confocal microscopic (40 X) studies of the *in-vivo* binding of FITC-custom peptides and scrambled peptides (50 µg/mL) to N2 *C. elegans* for 2h. ^*^p ≤ 0.05, a significant difference between the fluorescence intensity measured between FITC-TNP and HNP (50 µg/mL) to N2 *C. elegans*.

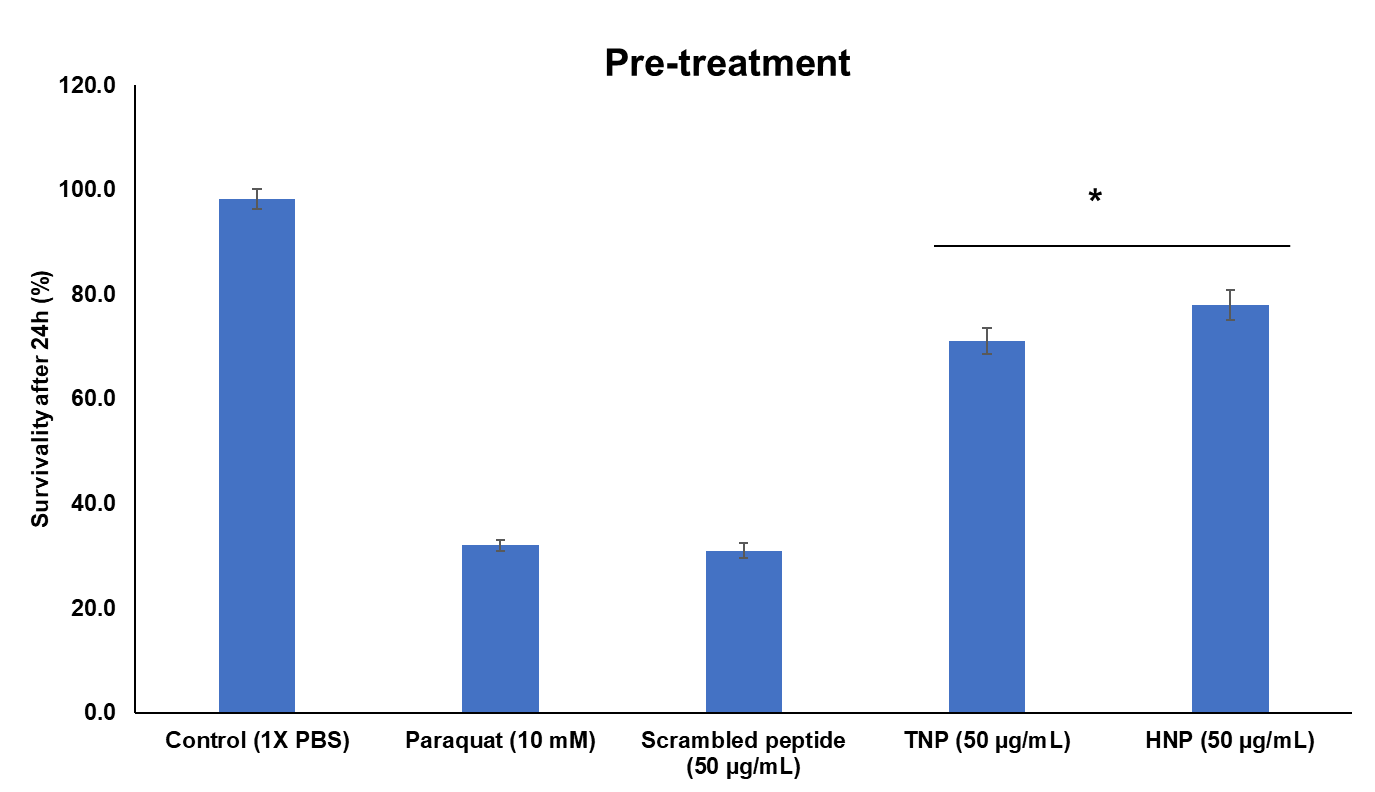

**Supplementary Fig S4.** Determination of the effect of the custom peptides and scrambled peptide on PT-induced death of wild type N2 strain of *C. elegans*. worms were pre-incubated with custom peptides/scrambled peptide (50 µg/mL) followed by the PT (10 mM) treatment. Percent survivavility was calculated against only paraquat-treated worms. ^*^p ≤ 0.05, a significant difference between only paraquat treated group and custom peptide (TNP, HNP).

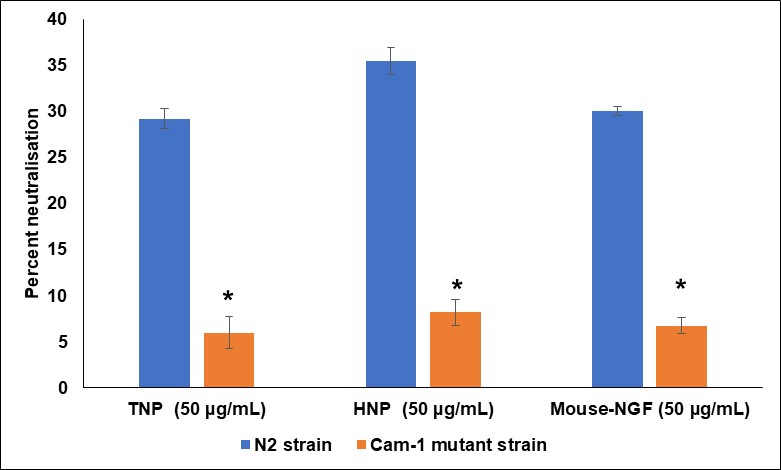

**Supplementary Fig S5.** Determination of the effect of the custom peptides and scrambled peptide on PT-induced death of wild-type N2 strain of *C. elegans*. worms were pre-incubated with custom peptides/scrambled peptides (50 µg/mL) followed by the PT (10 mM) treatment. Percent survivability was calculated against only paraquat-treated worms. ^*^p < 0.05, a significant difference between only paraquat treated group and custom peptide (TNP, HNP).

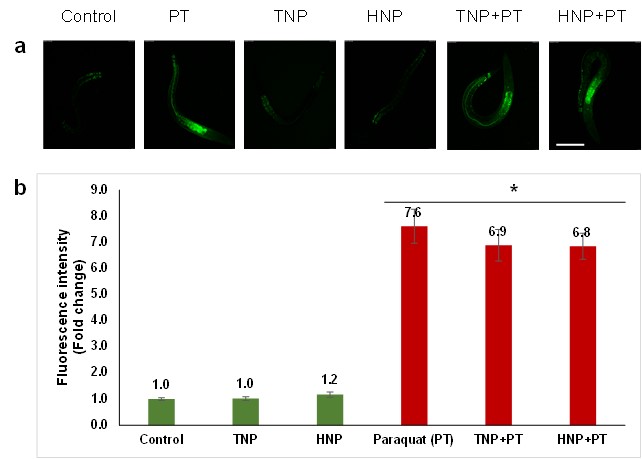

**Supplementary Fig S6.** Determination of PT-induced intracellular ROS generation and custom peptides pre-treatment could not restore the ROS generation in the cam-1 mutant strain of *C. elegans.* The ROS generation was determined by using an H_2_DCFDA fluorescence probe. **(a)** confocal microscope images of nematodes expressing ROS. The scale bar indicates the length as 100 µm. **(b)** Bar graph representing dosimetry analysis of confocal images to quantitate the intracellular ROS generation.

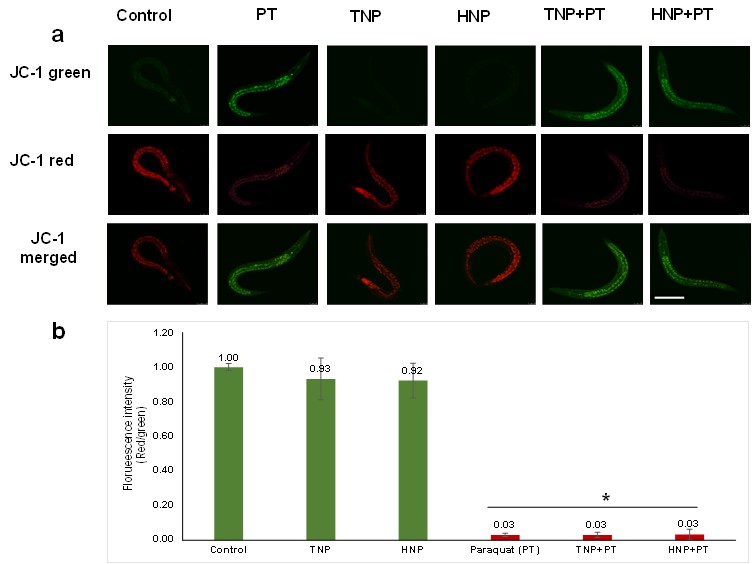

**Supplementary Fig S7. (a)** Confocal images of cam-1 mutant strain of *C. elegans* showing restoration of PT-induced disruption of mitochondrial membrane potential (MMP) of *C. elegans* pre-treated with custom peptides. The scale bar indicates the length as 100 µm**.** The scale bar indicates the size as 100 µm**.** The PT-treated (10 mM) N2 wild-type strain of nematodes pre-treated with or without custom peptide (50 µg/mL) was observed to measure the red/green fluorescence intensity ratio by JC-1 staining. **(b)** Bar diagram representing the red/green fluorescence intensity ratio quantified using Image J software. ^*^p ≤ 0.05, a significant difference between untreated (control)/only custom peptide treated group and PT/custom peptide and PT-treated group of cam-1 mutant worms.

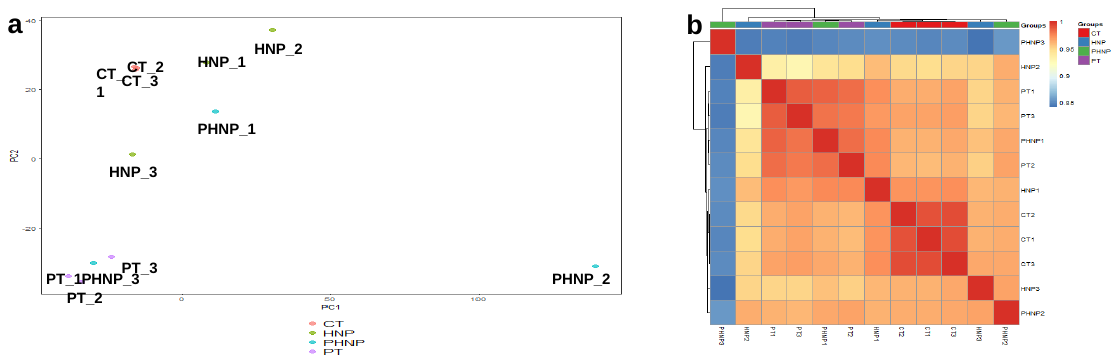

**Supplementary Fig S8. (a)** PCA score plot showing the gene expression variability between the groups of *C. elegans* and within the biological replicates. **(b)** Correlation plot showing a correlation between treated groups of *C. elegans*. CT: untreated worms, PT: PT treated worms, PHNP: custom peptide HNP pre-treatment followed by PT treatment, HNP: custom peptide HNP treated worms. Plots showing differential expression of genes in PT-treated *C. elegans* and their restoration with peptide (HNP) pre-treatment.

**Supplementary Fig S9.** The effect of the custom peptides (TNP: HNP:: 1:1) treatment on histological changes in the tissues of Swiss albino mice. The H and E staining was employed to observe any morphological changes in the tissues compared to those of the control (untreated). Light microscopic observation of a) Brain, b) Heart, c) Kidney, d) Liver, e) Lung, f) Ovary, and g) Testis for control and treated groups. Bar-100µM.

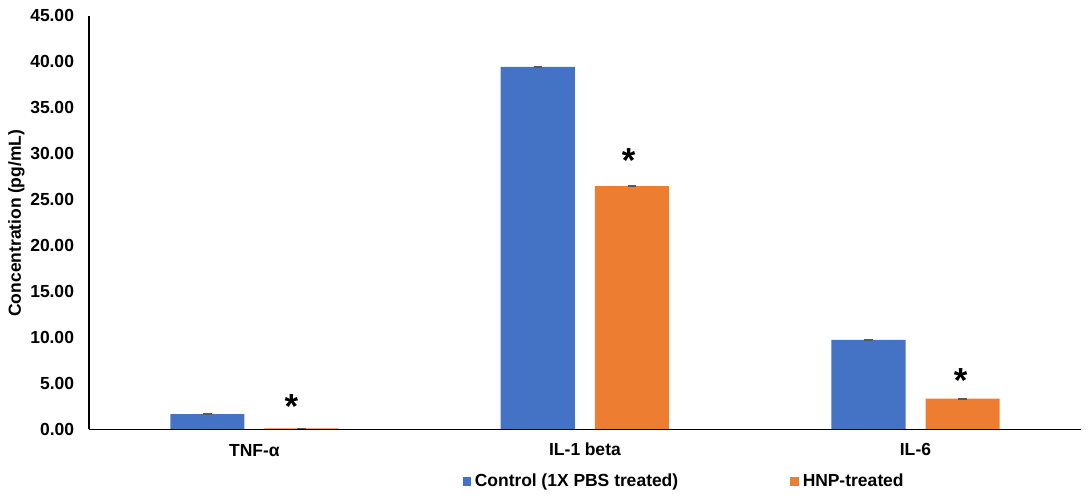

**Supplementary Fig S10**. Determination of the concentration of proinflammatory cytokines in control (1X PBS treated)/ custom peptides-treated (10 mg/kg) group of mice plasma by Quantikine HS ELISA Kit. ^*^ (p ≤ 0.05) a significant difference between control (1X PBS-treated) and custom peptide (HNP)-treated *C. elegans*. Values are means ± SD of triplicate determinations.

**Supplementary Table S1:** Primer sets used for qRT-PCR analysis.

| **Sequence name** | **Sequence** | **No. of Bases** |
| --- | --- | --- |
| sek-1_F | GCCGATGGAAAGTGGTTTTA | 20 |
| sek-1_R | TAAACGGCATCGCCAATAAT | 20 |
| pmk-1_F | CCGACTCCACGAGAAGGATA | 20 |
| pmk-1_R | AGCGAGTACATTCAGCAGCA | 20 |
| skn-1b_F | CTCTCTTCTGGCATCCTCTACCA | 23 |
| skn-1b_R | CCGACTCCACGAGAAGGATA | 20 |
| pmk-1_F | AGCGAGTACATTCAGCAGCA | 20 |
| pmk-1_R | TTCTTGGATTCTTCTTCTTGTTCGT | 25 |
| act-1_F | GCTGGACGTGATCTTACTGATTACC | 25 |
| act-1_R | GTAGCAGAGCTTCTCCTTGATGTC | 24 |
| sod-1_F | CGTAGGCGATCTAGGAAATGTG | 22 |
| sod-1_R | AACAACCATAGATCGGCCAACG | 22 |
| sod-3_F | TTCAAAGGAGCTGATGGACACT | 22 |
| sod-3_R | AAGTGGGACCATTCCTTCCAA | 21 |
| trx-1_F | TCCAACACTTTTTGACGCAG | 20 |
| trx-1_R | CAAGATGATGCCGACTTTCA | 20 |
| ser-1_F | AAGAGCCAGTCGCCAGAAC | 19 |
| ser-1_R | GTGGTTGATGCCTCTGTCGT | 20 |
| hsp-16.1_F | GCAGAGGCTCTCCATCTGAA | 20 |
| hsp-16.1_R | GCTTGAACTGCGAGACATTG | 20 |
| hsp-16.2_F | CTATTTCCGTCCAGCTCAAC | 20 |
| hsp-16.2_R | TTTGTTCAACGGGCGCTTGC | 20 |
| hsp-60_F | CTATGGGCCCAAAAGGAAGAAACGTG | 26 |
| hsp-60_R | GGATTTCGCGACGGTGACTCCGTCC | 25 |
| hsp-70_F | GAAAGGTTGAAATCCTCGCGAACTC | 25 |
| hsp-70_R | TCCGGATTACGAGCGGCTTGATCTT | 25 |
| ctl-1_F | CGGATACCGTACTCGTGATGA | 21 |
| ctl-1_R | CCAAACAGCCACCCAAATCA | 20 |
| ctl-2_F | TCCGTGACCCTATCCACTTC | 20 |
| ctl-2_R | TGGGATCCGTATCCATTCAT | 20 |
| gst-4_F | GATGCTCGTGCTCTTGCTG | 19 |
| gst-4_R | CCGAATTGTTCTCCATCGAC | 20 |
| gst-6_F | GGACAAGACTTCGAGGACAAC | 21 |
| gst-6_R | AACTGACGAGCCAAGTAACG | 20 |
| gst-10_F | AAGAGATTGTGCAGACTGGAG | 21 |
| gst-10_R  | AGAACATGTCGAGGAAGGTTG | 21 |
| ced-3_F | ACGGGAGATCGTGAAAGC | 18 |
| ced-3_R | AGAGTTGGCGGATGAAGG | 18 |
| ced-4_F | AGTCACTCGCAATGGCTCT | 19 |
| ced-4_R | GCTGATGAACGACGGAAT | 18 |
| ced-9_F | AAAGGCACAGAGCCCACC | 18 |
| ced-9_R | CGTTCCCATAACTCGCATC | 19 |

**Supplementary Table S2:** Comparison of the fold changes in differential expression of proteins in paraquat-treated *C. elegans* determined by proteomic analysis.

| **(a) List of upregulated proteins in paraquat-treated worms** **compared to untreated (control) worms and their downregulation in worms pre-treated with peptide HNP.** | | | | |
| --- | --- | --- | --- | --- |
| **Accession No** | **Paraquat treatment (fold change in expression compared to control)** | **HNP pre-treatment (fold change in expression compared to control)** | **Pathway name** | **Description** |
| sp\|P19974\|CYC21_CAEEL | 1.72 | 0.70 | Apoptotic pathway | Cytochrome c 2.1 OS=*Caenorhabditis elegans* OX=6239 GN=cyc-2.1 PE=1 SV=2 |
| G5EC98\|G5EC98_CAEEL | 1.62 | 0.60 | De novo pyrimidine ribonucleotides biosynthesis | CTP synthase OS=*Caenorhabditis elegans* OX=6239 GN=ctps-1 PE=1 SV=1 |
| sp\|O17071\|PRS10_CAEEL | 1.39 | 0.98 | Ubiquitin proteasome pathway | Probable 26S proteasome regulatory subunit 10B OS=*Caenorhabditis elegans* OX=6239 GN=rpt-4 PE=1 SV=2 |
| sp\|G5EEI4\|ASP1_CAEEL | 2.07 | 0.40 | Programmed cell death | Aspartic protease 1 OS=*Caenorhabditis elegans* OX=6239 GN=asp-1 PE=1 SV=1 |
| Q965Q1\|Q965Q1_CAEEL | 1.64 | 0.59 | Stress response | 10 kDa heat shock protein, mitochondrial OS=*Caenorhabditis elegans* OX=6239 GN=CELE_Y22D7AL.10 PE=1 SV=1 |
| P34707 (SKN1_CAEEL) | 1.29 | 0.75 | Neuronal cell death/ MAPK pathway | Protein skinhead-1 OS=*Caenorhabditis elegans* OX=6239 |
| O61667 (EGL1_CAEEL) | 1.97 | 0.54 | Programmed cell death | Programmed cell death activator egl-1 OS=*Caenorhabditis elegans* OX=6239 |

| **(b) List of downregulated proteins in paraquat-treated worms compared to untreated (control) worms and their upregulation in worms pre-treated with peptide HNP.** | | | | |
| --- | --- | --- | --- | --- |
| **Accession No.** | **Paraquat treatment (fold change in expression compared to control)** | **HNP pre-treatment, (fold change in expression compared to control)** | **Pathway name** | **Description** |
| sp\|P46561\|ATPB_CAEEL | 0.72 | 1.49 | ATP synthesis | ATP synthase subunit beta, mitochondrial OS=*Caenorhabditis elegans* OX=6239 GN=atp-2 PE=1 SV=2 |
| sp\|Q18688\|HSP90_CAEEL | 0.76 | 1.93 | Stress response pathway | Heat shock protein 90 OS=*Caenorhabditis elegans* OX=6239 GN=daf-21 PE=1 SV=1 |
| Q23682 (GCY5_CAEEL) | 0.70 | 1.73 | Chemosensory pathway | Receptor-type guanylate cyclase gcy-5 OS=*Caenorhabditis elegans* OX=6239 |
| A8XL75 (A8XL75_CAEBR) | 0.75 | 2.93 | Chemosensory pathway | Protein CBR-SRE-4 OS=*Caenorhabditis elegans* OX=6239 |
| sp\|P98080\|UCR1_CAEEL | 0.74 | 1.56 | ATP synthesis | Cytochrome b-c1 complex subunit 1, mitochondrial OS=*Caenorhabditis elegans* OX=6239 GN=ucr-1 PE=3 SV=2 |
| sp\|P11141\|HSP6_CAEEL | 0.71 | 1.35 | Stress response pathway | Heat shock protein hsp-6 OS=*Caenorhabditis elegans* OX=6239 GN=hsp-6 PE=1 SV=2 |
| sp\|Q05036\|HS110_CAEEL | 0.70 | 1.38 | Stress response pathway | Heat shock protein 110 OS=*Caenorhabditis elegans* OX=6239 GN=hsp-110 PE=3 SV=1 |
| sp\|Q18115\|PSMD1_CAEEL | 0.78 | 1.54 | Ubiquitin proteasome pathway | 26S proteasome non-ATPase regulatory subunit 1 OS=*Caenorhabditis elegans* OX=6239 GN=rpn-2 PE=3 SV=4 |
| O45445 (O45445_CAEEL) | 0.78 | 2.0 | Innate immune response | C-type lectin OS=*Caenorhabditis elegans* OX=6239 |
| sp\|P02513\|HSP17_CAEEL | 0.71 | 1.65 | Stress response pathway | Heat shock protein Hsp-16.48/Hsp-16.49 OS=*Caenorhabditis elegans* OX=6239 GN=hsp-16.48 PE=2 SV=1 |
| sp\|P10299\|GSTP1_CAEEL | 1.19 | 1.36 | Antioxidant pathway | Glutathione S-transferase P OS=*Caenorhabditis elegans* OX=6239 GN=gst-1 PE=1 SV=1 |
| sp\|Q09607\|GST36_CAEEL | 0.71 | 2.20 | Antioxidant pathway | Probable glutathione S-transferase gst-36 OS=*Caenorhabditis elegans* OX=6239 GN=gst-36 PE=3 SV=2 |
| sp\|P34697\|SODC_CAEEL | 0.54 | 1.38 | Antioxidant pathway | Superoxide dismutase [Cu-Zn] OS=*Caenorhabditis elegans* OX=6239 GN=sod-1 PE=1 SV=2 |
| sp\|P49632\|RL40_CAEEL | 0.52 | 1.40 | Ubiquitin proteasome pathway | Ubiquitin-60S ribosomal protein L40 OS=*Caenorhabditis elegans* OX=6239 GN=ubq-2 PE=3 SV=2 |
| Q21233\|Q21233_CAEEL | 0.62 | 1.44 | Electron transport chain | NADH dehydrogenase [ubiquinone] 1 alpha subcomplex subunit 10, mitochondrial OS=*Caenorhabditis elegans* OX=6239 GN=nuo-4 PE=1 SV=1 |
| Q17512\|Q17512_CAEEL | 0.76 | 1.65 | Electron transport chain | NADH Ubiquinone oxidoreductase Fe-S protein OS=*Caenorhabditis elegans* OX=6239 GN=nduf-11 PE=1 SV=1 |
| Q03561 (NLP36_CAEEL) | 0.60 | 1.66 | Neuronal development | Neuropeptide-like peptide 36 OS=*Caenorhabditis elegans* OX=6239 |
| Q86NC2\|Q86NC2_CAEEL | 0.75 | 1.42 | Electron transport chain | NADH dehydrogenase [ubiquinone] iron-sulfur protein 3, mitochondrial OS=*Caenorhabditis elegans* OX=6239 GN=nuo-2 PE=1 SV=1 |
| Q93727 (SPTF1_CAEEL) | 0.73 | 1.55 | Neuronal differentiation | C2H2-type domain-containing protein OS=*Caenorhabditis elegans* OX=6239 |
| Q93934\|Q93934_CAEEL | 0.79 | 1.65 | Autophagy | Adenosine kinase OS=*Caenorhabditis elegans* OX=6239 GN=adk-1 PE=1 SV=1 |
| Q9U329\|Q9U329_CAEEL | 0.74 | 1.65 | Oxidative phosphorylation | Cytochrome c oxidase subunit 4 OS=*Caenorhabditis elegans* OX=6239 GN=cox-4 PE=1 SV=1 |

| **(c) List of commonly expressed proteins between paraquat-treated worms and peptide HNP pre-treatment worms** | | |
| --- | --- | --- |
| **Accession No** | **Pathway name** | **Description** |
| Q966C7\|Q966C7_CAEEL | Pentose phosphate pathway | Transaldolase OS=*Caenorhabditis elegans* OX=6239 GN=tald-1 PE=1 SV=1 |
| sp\|H2KYQ5\|GYG1_CAEEL | Glycogen biosynthesis | Glycogenin-1 OS=*Caenorhabditis elegans* OX=6239 GN=gyg-1 PE=1 SV=1 |
| sp\|H2KYE0\|NRA4_CAEEL | Neuronal development | Nicotinic receptor-associated protein 4 OS=*Caenorhabditis elegans* OX=6239 GN=nra-4 PE=1 SV=1 |
| Q9N5S7\|Q9N5S7_CAEEL | Antioxidant pathway | Thioredoxin Domain Containing protein homolog OS=*Caenorhabditis elegans* OX=6239 GN=txdc-12.2 PE=1 SV=1 |
| sp\|Q9N4X8\|GSTPA_CAEEL | Antioxidant pathway | Glutathione S-transferase P 10 OS=*Caenorhabditis elegans* OX=6239 GN=gst-10 PE=1 SV=3 |
| G5EC71\|G5EC71_CAEEL | Antioxidant pathway | Glutathione S-Transferase OS=*Caenorhabditis elegans* OX=6239 GN=gst-20 PE=1 SV=1 |
| Q09590\|Q09590_CAEEL | Oxidoreductase | NADPH--cytochrome P450 reductase OS=*Caenorhabditis elegans* OX=6239 GN=emb-8 PE=1 SV=1 |
| sp\|Q11190\|ETFD_CAEEL | Electron transport chain | Electron transfer flavoprotein-ubiquinone oxidoreductase, mitochondrial OS=*Caenorhabditis elegans* OX=6239 GN=let-721 PE=3 SV=2 |
| sp\|G5ECU1\|SKR1_CAEEL | Neurogenesis | Skp1-related protein OS=*Caenorhabditis elegans* OX=6239 GN=skr-1 PE=1 SV=1 |

**Supplementary Table S3:** Some biochemical properties of serum of control and custom peptides (TNP and HNP; 1:1)-treated (10 mg/kg) mice after 24 hours of *i.v*. injection. Values are mean ± SD of 6 mice. There was no significant difference in values (p ≥ 0.05) between control and custom peptides-treated groups of mice.

|  | **Values** | |
| --- | --- | --- |
| **Parameters (Unit)** | **Control** | **Custom peptides (TNP and HNP)-treated** |
| Glucose (mg/dL) | 101.10 ± 1.27 | 160.0 ± 1.50 |
| Billirubin (Direct) (mg/dL) | 0.03 ± 0.00 | 0.08 ± 0.00 |
| Billirubin (Total) (mg/dL) | 0.05 ± 0.00 | 0.1 ± 0.00 |
| BUN (mg/dL) | 8 ± 0.05 | 10 ± 0.10 |
| Creatinine(mg/dL) | 0.18 ± 0.20 | 0.17 ± 0.20 |
| Albumin (mg/dL) | 1.98 ± 0.03 | 2.1 ± 0.05 |
| ALKP(U/L) | 81.85 ± 1.63 | 132 ± 1.70 |
| SGPT(U/L) | 33 ± 0.78 | 36 ± 0.70 |
| SGOT(U/L) | 62 ± 1.10 | 64 ± 1.13 |
| Cholesterol (mg/dL) | 66 ± 2.5 | 70 ± 3.0 |
